# STExplorer: Navigating the Micro-Geography of Spatial Omics Data

**DOI:** 10.1101/2025.01.17.633539

**Authors:** Eleftherios Zormpas, Nikolaos I. Vlachogiannis, Anastasia Resteu, Adrienne Unsworth, Simon Tual-Chalot, Birthe Dorgau, Rachel Queen, Majlinda Lako, Dina Tiniakos, Alexis Comber, Quentin M. Anstee, Antonis Giakountis, Simon J. Cockell

## Abstract

Spatial transcriptomics (ST) has the potential to provide unprecedented insights into gene expression across tissue architecture, but existing analytical methods often overlook the full complexity of the spatial dimension. We present **STExplorer**, an R package that adapts well-established computational geography (CG) methods to explore the micro-geography of spatial omics data. By incorporating techniques like Geographically Weighted Principal Component Analysis (GWPCA), Fuzzy Geographically Weighted Clustering (FGWC), Geographically Weighted Regression (GWR), and analyses of observation Spatial Autocorrelation (SA), STExplorer enables the uncovering of spatially resolved patterns that capture the spatial heterogeneity of biological data.

STExplorer provides a complete suite of functions for spatial analyses and visualisations, supporting deeper biological understanding and inference. Built on the Bioconductor ecosystem, the package integrates with SpatialFeatureExperiment objects, ensuring compatibility with existing pipelines. It includes preprocessing capabilities such as data import, quality control, gene count normalisation, and variable gene selection, alongside tools for downstream analysis and detailed visualisations that quantify and map spatial heterogeneity and relationships.

We demonstrate the utility of STExplorer through applications to spatial transcriptomics datasets, revealing that spatially varying gene expression and relationships are often masked by standard analyses. By bridging bioinformatics and CG, STExplorer provides a novel and informed approach to spatial transcriptomics analysis, with robust tools to address spatial heterogeneity and its associated underlying biology, thereby advancing our understanding of complex tissue biology without reinventing the wheel.

## Main

Since the 1960s, when *in situ* hybridisation (ISH) was first established, a series of technological advancements have gradually enhanced our understanding of tissue function by examining the spatial landscape, molecular biology, and the spatial interactions of cells that comprise tissues[1]. Methodological developments are moving towards the ability to capture whole-transcriptome molecular information at a single-cell level while retaining spatial information. Initially, modern ISH-based technologies like single-molecule fluorescence *in situ* hybridisation (smFISH)[2] and MERFISH[3], or others like STARmap[4] and GeoMx[5] provided the means to measure gene expression at single-cell level in a spatially-aware manner. Although presently these methods do not scale beyond just a few thousand genes with GeoMx providing the “Whole Transcriptome Atlas” offering 18,000 targets for human tissues[5]. This does not allow for an unbiased, hypothesis-free examination of the spatial organisation of gene expression. To this end, single-cell RNA sequencing (scRNA-Seq) has been an invaluable technology for generating unbiased, hypothesis-free data at single-cell resolution, without relying on pre-defined target-gene probe panels. Many approaches exist for scRNA-seq[6], all of which share a common step of dissociating the cells leading to loss of spatial context. Since the publication of the first sequencing-based spatial transcriptomics (ST) method[7], we are now able to assay gene expression both spatially and comprehensively (whole transcriptome). In less than a decade, a multitude of sequencing-based ST assays have been developed such as 10X Genomics Visium, Slide-seq/V2[8, 9], Stereo-seq[10], and Visium HD, which offer increasing levels of resolution ranging from 2-10 cells (Visium) to sub-cellular level (Stereo-seq/Visium HD).

Sequencing-based ST approaches generate spatially oriented expression data where the digital read count and their spatial properties are entangled. This intricate relationship requires a different approach to analysis. Overlooking the spatial properties while investigating gene expression can compromise the integrity of the analysis, potentially overlooking crucial insights. High throughput spatial datasets are relatively new in biology. However, other disciplines have been dealing with such data for decades, continuously developing methods and approaches to use them appropriately while also accurately describing their characteristics. Research in the geographic sciences has identified three key aspects of spatially resolved datasets that are crucial to understand and incorporate in spatial data analysis. Namely, these are the Modifiable Areal Unit Problem (MAUP)[11], Spatial Autocorrelation (SA)[12], and Spatial Heterogeneity (SH)[13]. Each of these features is universal to spatially oriented data and has been shown to manifest in ST datasets[14]. Firstly, the MAUP highlights the impact of spatial data aggregation scale on statistical relationships and analytical interpretations. Secondly, SA emphasises the tendency for nearby observations to exhibit similarities, challenging the assumption of observation independence typically applied in classical statistics. Lastly, SH, or spatial non-stationarity, acknowledges that local relationships between covariates in spatial data can differ from global relationships[14]. In recent years, there has been a surge of publications using ST technologies, and this number is steadily increasing. In many of these publications, spatial information is usually employed to predefined specific areas of interest. However, even a simple task such as predefining specific areas of interest can introduce statistical challenges associated with spatial aggregation, hinting at MAUP, SA and SH. Yet, the true value of spatial content emerges when we integrate observation coordinate information as an integral element of the data analysis[14].

Appropriate consideration of spatial data analysis issues, as studied extensively in disciplines such as geography and ecology[15–21], alongside the treatment of coordinates as a covariate, lead to a new type of spatial transcriptomics data analysis. Utilising the spatial dimension, this analysis attempts to further our understanding of how gene expression is organised in space[14]. For example, Computational Geography (CG) has a rich history of method refinement and development in analysing spatial patterns and interactions in spatial data[19, 22–25]. As a result, lifting existing knowledge from CG permits a cross-disciplinary and collaborative approach to spatial data analysis allowing us to build coherent foundations in exploring spatial biology without reinventing the wheel.

Here we introduce Spatial Transcriptomics Explorer (STExplorer) (https://github.com/LefterisZ/STExplorer), an R package providing a set of methods derived from the field of geography, modified to be applied in the micro-scale geography of the tissue. STExplorer is a suite of CG-inspired methods: Geographically Weighted Principal Components Analysis (GWPCA)[26], Fuzzy Geographically Weighted Clustering (FGWC)[27], Spatial Autocorrelation calculations (SA)[28–32], and Geographically Weighted Regression analysis (GWR)[13]. These geo-computational methods, modified and applied to ST data, can assist us in exploring biology locally rather than dataset-wide, revealing the process heterogeneity over space that needs to be accounted for in our analyses. The vision behind the conception and continuous development of STExplorer is simple: there are a host of considerations, methods, and tools in CG that are currently not being exploited in the field of ST[14]. CG tools for handling spatial data and spatial problems have long pedigrees and have been subject to several refinements. Consequently, these are well established with clear benefits to leveraging these tools into ST practice rather than “reinventing” or “rediscovering” them.

## Results

### STExplorer Overview

STExplorer is an R-based toolkit of CG methods modified to apply to bio-spatial data like ST. The toolkit currently implements four classes of analysis from computational geography: GWPCA, FGWC, GWR, and SA calculations (Fig. 1). As input, STExplorer accepts spatial transcriptomics data (e.g. 10X Visium). For the initial steps of the analysis, involving data loading, Quality Control (QC), gene count normalisation, and gene selection, STExplorer utilises pre-existing methods from the Bioconductor ecosystem. This allows compatibility with other ST data analysis pipelines operating within Bioconductor (Fig. 1A). Specifically, the data are first imported into SpatialFeatureExperiment (SFE) objects using the import methods from the *SpatialFeatureExperiment*[33] R package. The advantage of using the SpatialFeatureExperiment object class for STExplorer is that it already incorporates the standardised way to encode spatial vector data in geographical sciences namely *Simple Features for R* (SF)[34]. Additionally, the SF package allows us to manipulate space and generate and store geometries for neighbour selection and visualisations in a uniform and standardised way.

**Figure 1.**
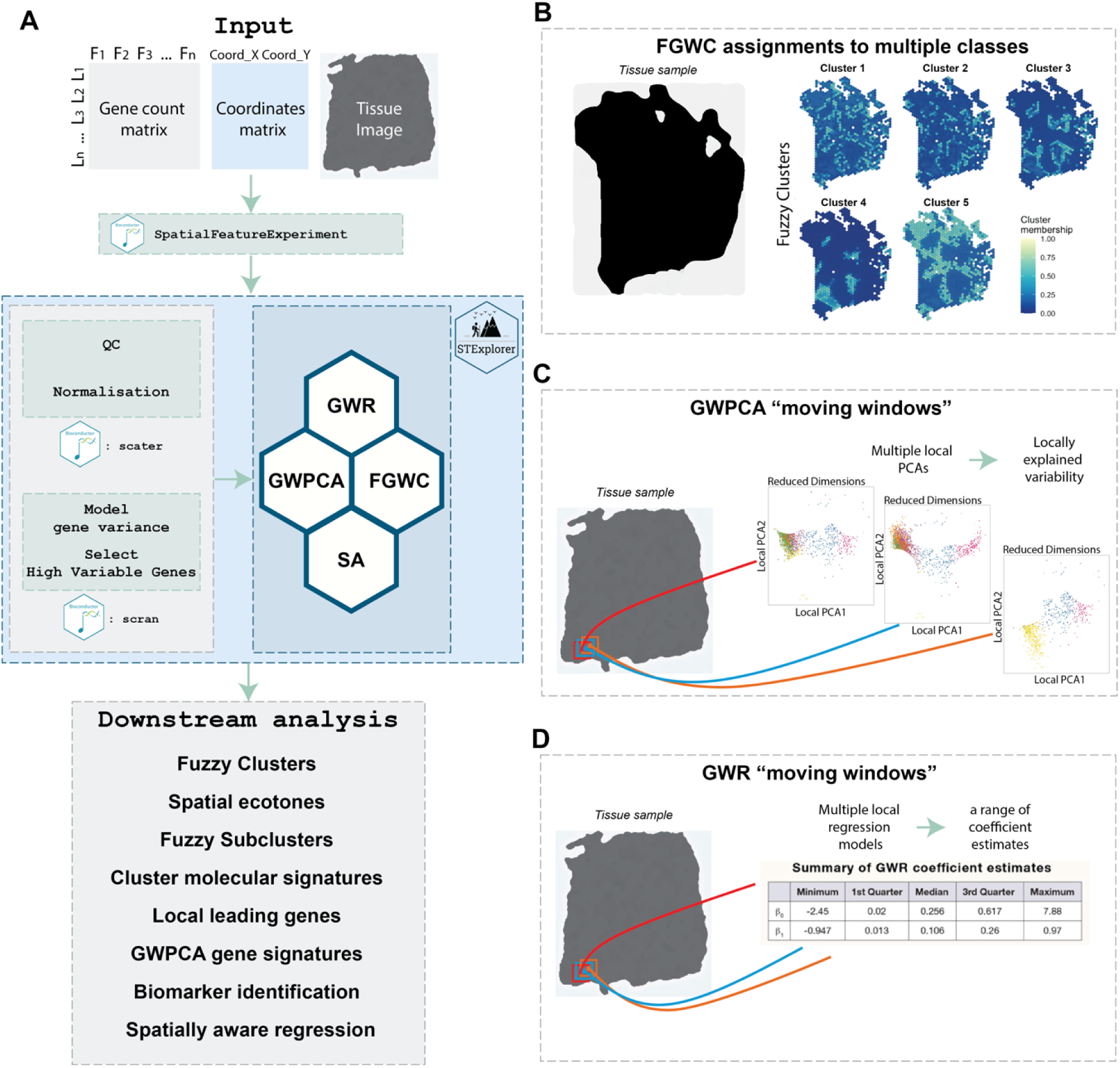
STExplorer overview. Overview of the pipeline and main components of STExplorer. **A.** STExplorer toolkit pipeline for the analysis of Spatial Transcriptomics data. The input is a gene count matrix, a coordinates matrix and an image (optional). The quality control (QC), normalisation, gene variance modelling, and high-variable genes (HVGs) selection are executed by the Bioconductor (http://bioconductor.org/) R packages scater and scran. In the next step, the FGWC, GWPCA, GWR and SA CG methods are utilised to perform a spatially-aware analysis of ST data. The output of those methods is then utilised further to extract biologically meaningful information from the data. **B.** FGWC is a supervised clustering method that assigns a membership to each class (cluster) for each location. Thus, the output is a series of class memberships maps rather than a single map of unique-class assignments. **C.** The moving windows logic of GWPCA. GWPCA calculates multiple localised PCAs while using distance weighted data subsets to determine the gene expression in the current window. The user sets the window size. GWPCA is complete once the moving window passes through every location. **D.** The moving windows logic of GWR. In essence, GWR calculates multiple spatially weighted regression models, one for each location. The result is a range of mappable coefficients for each neighbourhood.

Once the data are loaded, multiple QC metrics are added to the SFE object via functions from the *scater* R package[35] alongside some custom metrics and geometries required by the STExplorer to operate downstream. Gene count normalisation is also provided by the *scater* R package – using the default library size normalisation with *log2* transformation of gene counts. Nonetheless, the pipeline is compatible with other types of normalisation, such as SCTransform from the Seurat[36] package. Finally, STExplorer utilises *scran*[37] for modelling the gene variance and for selecting high variable genes (HVGs) (Fig. 1A). However, gene selection for downstream analysis can be executed by other approaches (e.g., identify spatially variable genes) or packages (i.e., Seurat) and fed back into STExplorer.

After completing the pre-processing steps, the STExplorer toolkit pipeline allows users to perform FGWC, GWPCA, and GWR analyses on the data while providing the means to allow proper utilisation of the outputs downstream (Fig. 1A). Geographically Weighted frameworks like FGWC, GWPCA, and GWR use moving window (kernel) based approaches to extract data subsets, from which local models, statistics or metrics are calculated. The observations in the subsets are weighted by their distance to the location being considered[38]. A critical consideration is the kernel bandwidth. This describes the size of the moving window (fixed distance), or the proportion of the data points used in each local model (adaptive distance bandwidth). In this way, the bandwidth defines the degree of smoothing in the outputs, and many GW models include approaches to determine bandwidths that optimise model error or fit.

FGWC is a spatial data analysis technique combining fuzzy clustering principles and geographically weighted modelling. The approach extends the traditional fuzzy clustering methodology[27] by integrating spatial heterogeneity, allowing clusters to adapt to local variations (see Methods). In traditional clustering, each data point belongs to a single cluster. In FGWC, a data point (in ST data, this can be a spot, a bin or a location) has partial membership in every cluster (Fig. 1B). Geographic weighting of the gene expression observations influence the FGWC process according to spatial proximity, resulting in a more nuanced clustering solution that reflects the inherent uncertainty of data categorisation and the varying characteristics of spatial datasets.

GWPCA is an extension of Principal Component Analysis (PCA), which effectively incorporates the concepts of spatial heterogeneity and spatial autocorrelation. GWPCA utilises a “moving windows” approach (Fig. 1C), allowing the analysis of spatially varying relationships in multivariate data by modifying PCA to account for the spatial distribution of observations (see Methods). GWPCA analyses local relationships between variables by computing principal components (PCs) that vary across different locations in the study area. This method provides insights into how patterns of variation in multivariate data change over space Traditional regression models (e.g., ordinary least squares (OLS)) typically assume that the relationships being modelled are consistent (stationary) across the entire study area, employing parameters that represent ’whole-map’ statistics[23, 39]. Yet, this assumption is often untrue as there might be intrinsic relationships over space, and thus, regression parameters must be adjusted in light of this[39]. GWR is an effective method in exploratory spatial data analysis to understand the relationship between variables over space and reveal “shifts” in relationship strength and/or direction in different areas. GWR follows a principle similar to that of GWPCA – the moving windows – and calculates a spatially weighted model (or spatially variable coefficient (SVC) model) in each neighbourhood (Fig. 1D). The SVC models generate coefficient estimates at each location for each predictor variable. As such, they allow the spatial heterogeneity in modelled relationships to be explored and mapped.

### FGWC analysis in prostate cancer

FGWC originates from Bezdek’s fuzzy c-means algorithm[40] and represents a different approach from absolute Boolean clustering by combining the concepts of partial class membership and spatial proximity. Instead of simply assigning each observation to a single class or cluster, FGWC determines the membership of each observation to each class. This method recognises that observations or areas may have partial characteristics (and thus membership) of multiple clusters [27].

FGWC applied to bio-spatial data assumes that biological clusters often lack hard boundaries, instead display ecotones at their borders[41], which reflect gradients between spatially adjacent classes (Supp. Fig. 1). A concept borrowed from ecology, an ecotone describes the transition zone between two biological communities, with variable widths depending on transition “smoothness.” On the micro level, biological ecotones may arise naturally or through experimental platforms (Supp. Fig. 1). Low-resolution platforms, like Visium, can create ecotones via overlapping cell zones, while high-resolution platforms, like Stereo-seq or Visium-HD, may introduce artificial ecotones by binning data. Even with single-cell resolution (e.g., Stereo-seq with cell segmentation), ecotones can appear as cellular mosaics within tissues or as cells transitioning between stages, naturally occurring within tumours as histologically diverse regions of malignant, immune, stromal, and epithelial cells.

To challenge ecotone identification through FGWC, we used a 10X Visium-SD histologically annotated prostate cancer dataset[42]. The patient biopsies represent tissue sections with diverse combinations of stroma, benign, inflamed and cancerous areas. FGWC successfully isolated fuzzy clusters that accurately reflect the underlying histology (Fig. 2A). Subsequently we can select the cluster with the highest percentage of membership in each location (Fig. 2B) and superimpose it as a cluster map on the tissue section (Fig. 2A, Suppl. Fig. 2A). If we compare the highest membership cluster map with the histological annotation, it becomes clear that the FGWC results are in broad agreement with the histopathological assessment of the tissue section, building towards some interesting conclusions. For example, the histological annotation of section H2_5 separates the cancerous lesion (upper part of tissue section) from transition and surrounding stroma areas (upper left side of tissue section), while FGWC assigns the whole upper region as cancerous with the partial inclusion of surrounding stroma and transition areas on the left (cluster 4), suggesting that at the molecular level cancer is already established in these areas (Fig. 2A). Importantly, the same stroma/transition regions are also represented in cluster 2, yet with lower membership scores compared to the remaining stroma spots, defining a *bona fide* case of ecotone identification. FGWC independently assigns another stroma/transition area as a separate cluster (3), suggesting that the stroma is far more heterogeneous at the molecular level than the existing histological annotation.

**Figure 2.**
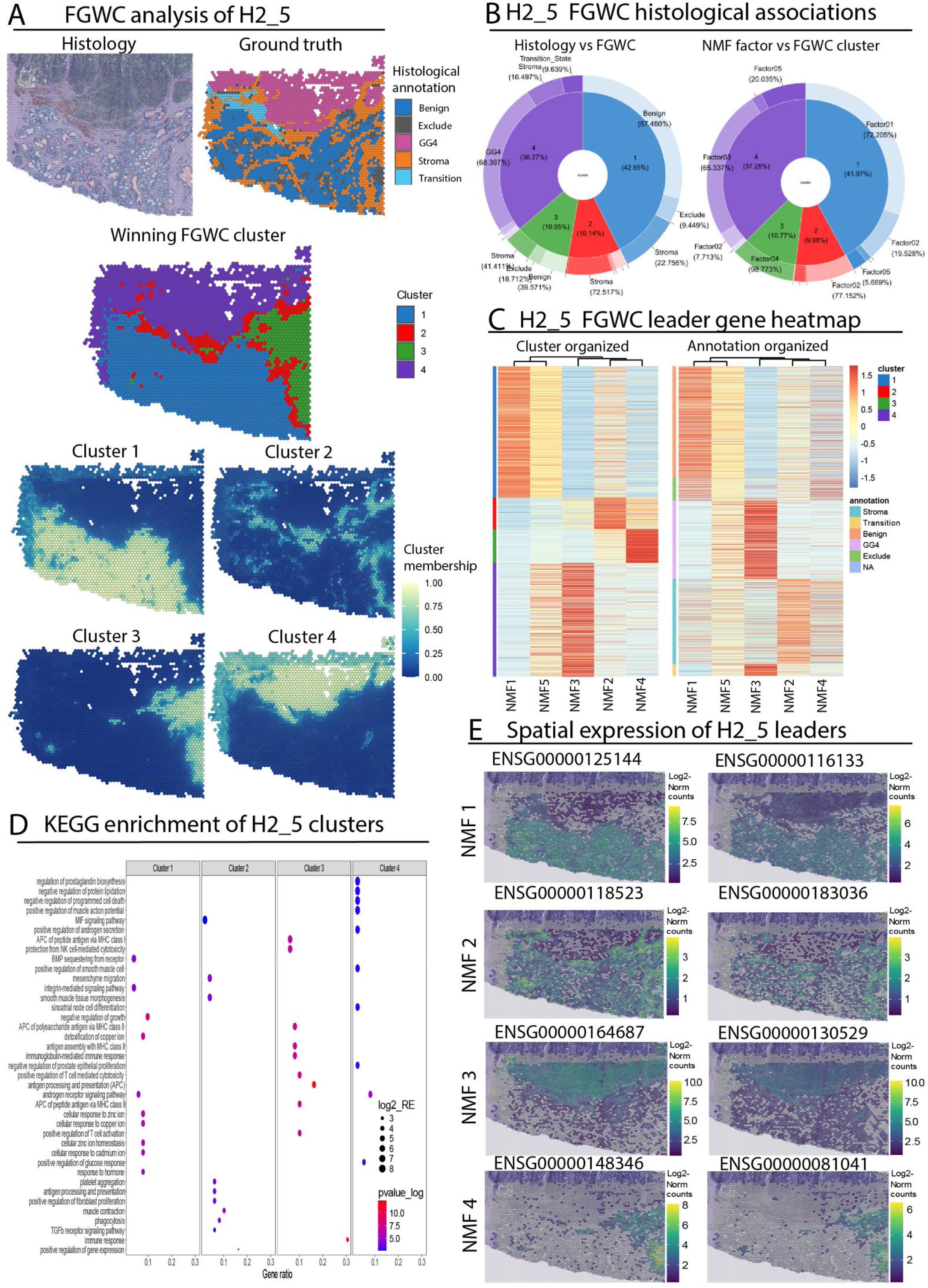
**Fuzzy Geographical Weighted Clustering analysis of prostate section H2_5**. **A.** Comparison of FGWC clusters with histological annotation for prostate tumour section H2_5, composed mainly of GG4 cancerous (top) and benign stroma/epithelium (bottom). **B.** Pie-Donut plot illustrating the distribution of FGWC clusters per histological annotation (left) and NMF factors per FGWC cluster (right) for H2_5. **C.** Heatmap highlighting the leading genes per NMF factor organized according to FGWC clusters (left) or histological annotations (right) **D.** KEGG pathway enrichment analysis for the leading genes of each FGWC cluster **E.** Representative examples of spatial gene expression for selected NMF factors. ***Gene References:*** *ENSG00000125144* ⇔ *MT1G*[47, 48]*, ENSG0000011C133* ⇔ *DHCR24[4S], ENSG000001C4C87* ⇔ *FABP5*[50, 51]*, ENSG0000013052S* ⇔ *TRPM4*[52, 53]*, ENSG0000014834C* ⇔ *LCN2[54-5C], ENSG00000081041* ⇔ *CXCL2*[57, 58]

Within the same section, spots previously annotated as benign are organised as a single and well-defined cluster (1) at the border of the cancerous tissue. Importantly, a region of stroma cells, marked with cluster 2, insulates the GG4 cancerous spots (cluster 4) from benign epithelium (cluster 1). The independent identification of cluster 2 (stroma at the border of the tumour) could very well reflect the influence that the tumour may exert on stroma at the cellular and molecular level (e.g. stroma residing inflammatory CD8+ cells attacking the tumour). H1_5 is a section dominated by GG4 with a few spells of cancerous stroma (Suppl. Fig. 2A). FGWC performs well in separating non-cancerous stroma from section areas in which cancer is present (Suppl. Fig. 2A). In contrast, H1_4 is more heterogeneous with cancerous, stroma and benign spots intermingled. Again, FGWC accurately defines all annotations as separate clusters (Suppl. Fig. 2A, GG4 and GG2 are merged into the same cluster (3) here). Interestingly, cluster 2 exclusively marks benign epithelium close to cancerous areas, separating it from distant benign/stroma cells (clusters 4&5). These results highlight the agreement between FWGC and the underlying histology, supporting the use of FGWC for analysing both complex and histologically homogenous tissue sections.

### Identification of FGWC/NMF metagene signatures

The input for FGWC derives from non-Negative Matrix Factorisation (NMF) of selected highly variable genes. NMF uncovers a lower-dimensional representation of the gene expression data in each location as metagene signatures. This parts-based data representation is interpretable and useful in understanding the molecular functions that underpin each cluster. The NMF factors consist of gene sets in the data aligning with the most informative components of key locations. Using the integrated pie-doughnut plot functions of STExplorer, the user can identify which individual NMF factor (or combination thereof) is more prominent in each location, connecting the cluster of locations with specific low-dimensional components identified by the original NMF (Fig. 2B).

Consequently, the NMF results allow the STExplorer user to identify prioritised (based on NMF score) metagene leader signatures that mediate the underlying biological functions, shaping histologically informative FGWC clusters. Based on the NMF output, we can produce annotated heatmaps of the metagene signatures (NMF factors) contrasting the expression of their constituent genes against the histological annotation and unveiling information that is not obvious from the ground truth. For example, in section H2_5 there is not a 1:1 relationship between FGWC clusters and NMF factors. Despite this, some clusters have a strong relationship with a single NMF factor that defines the cluster (e.g. cluster 3, Fig. 2C). However, FGWC considers all factors giving rise to unique patterns that identify each cluster but are not always dependent on a single factor. These patterns are not equally obvious when the heatmap is organised according to the underlying histology, highlighting the analytical power of the NMF/FGWC analysis for complex tissues (Fig. 2C).

Given that metagene signatures represent diverse cellular states with distinct molecular features, we can subject them to functional enrichment analysis, providing mechanistic insights for their corresponding clusters. For example, KEGG pathway analysis of the metagene signature from cluster 1 of section H2_5 (benign epithelium) is enriched in functions that are typical for prostate epithelium, such as androgen receptor signalling and negative regulation of growth. This contrasts with cluster 4 (GG4) which is enriched in processes associated with the negative regulation of programmed cell death, prostaglandin biosynthesis and positive regulation of androgen secretion (Fig. 2D, Suppl. Table 1). On the other hand, cluster 2 (stroma) is enriched in functions associated with antigen processing and presentation, positive regulation of fibroblast proliferation, muscle contraction and TGFa receptor signalling. Interestingly, cluster 3 which corresponds to a distinct and well-defined stroma ecotone, is enriched for APC antigen presentation, T-cell mediated cytotoxicity, and antigen assembly (MHC class I & II), suggesting that these stroma spots could be enriched in CD8^+^ and CD4^+^ cells that mediate an active inflammation response (Fig. 2D). STExplorer also provides integrated functions that allow the user to map the expression of leading genes of interest for each NMF factor, which can be used to validate the results of the functional analysis (Fig. 2E).

The FGWC/NMF pipeline is equally effective in purely benign sections like V1_2 (Suppl. Fig. 2B-E). In this section, FGWC identifies five clusters, allowing for the examination of ecotones between stroma and benign areas, with and without inflammation (Suppl. Fig. 2D). These clusters are described at the molecular level by various NMF factors (Suppl. Fig. 2B). For instance, NMF factor 2 corresponds closely with stroma spots and Fuzzy cluster 3, while NMF factor 1 primarily represents benign epithelium and Fuzzy cluster 4. Cluster 4, although covering areas that mainly consist of stroma cells and characterised by NMF factor 1, shows spots on its outskirts (Suppl. Fig. 2D) with intermediate memberships to adjacent cluster 3, suggesting ecotones formed by a mixture of benign epithelial and stroma cells. Each NMF factor is described by a metagene signature[43]. These signatures can be used to isolate the highest scoring genes (Suppl. Fig. 2C) and visualise their pattern of spatial expression (Suppl. Fig. 2E). This reveals, for example, that cluster 1 and NMF factor 4 define a stroma-associated ecotone which expresses genes such as LTF (ENSG00000012223) that regulates the immune microenvironment of prostate cancer[44] and FOS (ENSG00000170345), the inactivation of which can promote prostate cancer progression[45]. Finally, many locations are annotated as “Exclude” between this factor (NMF4) and NMF5. These areas are not annotated by histopathology. Nonetheless, FGWC added them to a category with a certain probability percentage, suggesting that molecular insights driven by spatial transcriptome analysis can overcome the lack of histological input. These findings indicate that FGWC/NMF can detect transcriptional alterations both in uniform and heterogenous tissue sections, highlighting distinct ecotones for downstream analyses.

Additionally, to further test FGWC, we analysed an eight post-conception week developing eye sample[46] (Supp. Fig 3). The tissue layers in this sample are much thinner and cover fewer spots than in the prostate cancer sample. Furthermore, as this is a developmental sample the cell type and structure of the eye at this stage of development are less well-defined than adult tissue. This poses an excellent case for testing the fuzziness of FGWC. Despite these challenges, FGWC effectively identified regions of biological interest. FGWC identified 8 clusters (Supp. Fig. 3), which broadly correspond to the ground truth annotation[46], demonstrating its robustness in capturing meaningful biological and structural patterns in this challenging dataset.

### FGWC/NMF-derived metagene signature performance in prostate bulk RNA-seq

Since FGWC/NMF provides prioritised lists of fuzzy cluster-specific genes that mediate neoplastic and physiological functions, one can expect that these metagene signatures could have a clinical footprint in bulk RNA-seq data. To test this hypothesis, we challenged FGWC/NMF results beyond spatial analysis, subjecting 13 NMF-derived metagene signatures from all tissue sections (Suppl. Table 2) to bulk RNA-seq analysis using a combination of The Cancer Genome Atlas (TCGA) prostate adenocarcinoma dataset (PRAD) representing cancerous data and the Genotype-Tissue Expression (GTEx) equivalent, representing normal prostate expression. Each signature consists of the top 10 genes associated with cancerous, benign or stroma clusters based on NMF factor scores. We confirmed that all metagene signatures are differentially expressed in the prostate bulk RNA-seq dataset, following a deregulation pattern that fully aligns with their origins in the spatial prostate dataset (Suppl. Fig. 4A, Suppl. Table 3).

Additionally, ROC analysis confirmed that the NMF signatures show a significant diagnostic potential for prostate cancer, with elevated AUC performance being observed for both cancerous and benign signatures (Suppl. Fig. 4B, Suppl. Table 4). In most cases, metagene signature expression was significantly associated with tumour stage, Gleason primary score or lymph node invasion, supporting their robust performance in bulk RNA- seq data and hinting towards their prognostic significance (Suppl. Fig. 4C). Collectively, FGWC coupled to NMF provides an accurate way of utilising the underlying transcriptional variability for the analysis of spatial heterogeneity. Moreover, despite inter-patient variability, NMF-derived metagene signatures behave robustly in bulk RNA- seq data and can be utilised as diagnostic/prognostic biomarkers. Another crucial advantage of the integrated FGWC/NMF approach is the opportunity to analyse complex sections without histological annotation. In such a scenario, FGWC clusters can accurately substitute the missing ground truth, while their associated NMF metagenes can assign molecular functions to the corresponding areas within the tissue section, helping the user to annotate the slice through the use of previously established tools such as the molecular anchoring functions of the Seurat pipeline. To the best of our knowledge, STExplorer is the only package to date that provides the user with such a framework to explore spatial clusters.

### GWPCA GSEA in prostate cancer

Global components of variability can shape gene expression patterns across an ST dataset. However, these global components cannot always explain all sources of locally observed variability, leaving a substantial portion unexplained and unexplored. STExplorer offers a second analytical axis, GWPCA, which facilitates the investigation of local components of variability in complex tissue structures. Users can apply the GWPCA functions directly on their spatial data or in combination with an FGWC/NMF analysis, the results of which can complement or even guide GWPCA, especially in heterogenous tissues that lack histological annotation. GWPCA provides the user with the top-ranking gene or prioritised lists of multiple leading genes. These can be further analysed as molecular sources contributing to the locally observed variability.

Cancer biopsies frequently comprise a dynamic and highly variable microenvironment, representing an ideal biological source for challenging GWPCA performance. For example, in the prostate cancerous section H1_4, non-cancerous stroma and benign epithelium co-exist with GG2 and GG4 cancerous areas (Fig. 3A). GWPCA reveals the existence of several leading genes even within histologically uniform areas, such as the non-cancerous stroma and/or the benign epithelium. Similarly, molecular complexity is also present in homogenous cancerous sections such as H1_5 or even in benign biopsies like V1_2 (Suppl. Fig. 5A). In large areas of both sections, a single gene can act as a leader, in contrast to neighbouring spots in which multiple genes share leadership roles in a dynamic manner, revealing a molecular mosaic that governs an otherwise homozygous histological annotation. As a consequence, the percentage of total variation (PTV) explained per location varies significantly. In the case of the cancerous section H1_4, the first ten PCs can, on average, explain around 50% of the variability in any particular location (Fig. 3B). On the contrary, for the benign slice V1_2, the first ten PCs can explain, on average, around 70% of the variability in a certain location. GWPCA represents a second, independent analytical axis of STExplorer that applies principal component analysis individually across multiple neighbourhoods of the tissue section, compressing and assigning transcriptional variability into spatially weighted components. In alignment with the NMF score strategy, the gene loadings of each GWP component can be used to select prioritised lists of leading genes for downstream functional analysis (Suppl. Tables 5 and 6).

**Figure 3.**
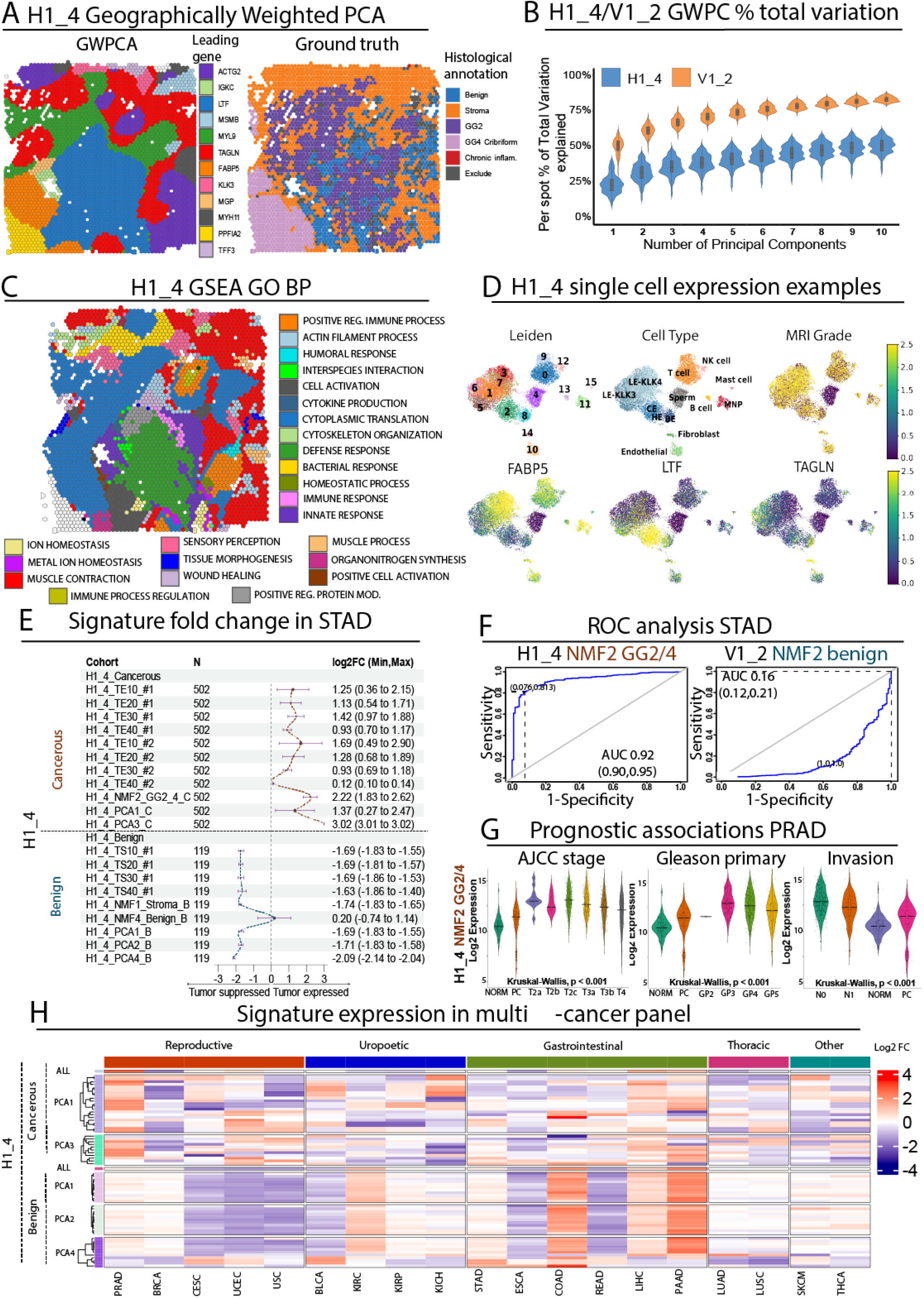
GWPCA analysis of spatial transcriptome data in prostate cancer sections. A. GWPCA maps visualizing the top leading genes in the cancerous prostate section H1_4. Ground truth representing histological annotation is also shown as a section map for comparison. **B.** Cumulative percentage of total variation (PTV) that is explained by the first 10 GWPCA Principal Components for H1_4 and V1_2 samples. Each violin plot consists of the PTV values of each individual spot in a sample. **C.** GSEA map summarizing the GO BP functional enrichment results for section H1_4 **D.** scRNA-seq UMAPs summarizing the expression of three representative leading genes from the prostate section H1_4. Leiden clusters along with cell type annotations and MRI grade are also shown for comparison **E.** Forest plot summarizing the fold change of twenty FGWC and GWPCA metagene signatures for section H1_4 in TCGA-PRAD data. Expression change is shown as log2FC along with min and max changes in parenthesis. The vertical dashed line corresponds to absence of differential expression, red curves correspond to upregulation in prostate tumour biopsies while blue curves correspond to upregulation in normal (GTEx) prostate biopsies. TS: Tumour-suppressed, TE: Tumour-expressed, TE10-40: GWPCA1 top 10-40 genes, PCA1-4: GWPCA1-4 top10 genes. **F.** Representative ROC-AUC analysis for prostate tumour diagnosis of one cancerous (left) and one benign (right) NMF meta-signature from H1_4. Average AUC with 95% CI are shown in parenthesis along with sensitivity and specificity performance. **G.** Violin plots illustrating the expression of the NMF2 GG2/4 cancerous meta-signature from section H1_4 in TCGA-PRAD biopsies stratified according to AJCC tumour stage (left), primary Gleason score (centre) or AJCC lymph node invasion (right). Kruskal-Wallis corresponds to statistical comparisons against the expression in the normal biopsies. **H.** Heatmap summarizing the fold change of all H1_4 GWPCA meta-signatures across a multi-cancer TCGA panel, shown as z-scores. Red corresponds to average log2 fold chance increase, blue corresponds to average log2 fold change decrease of each signature in the corresponding cancer type. Tumour types are organized according to system origins as indicated with the top-coloured bar. Cancer names are shown at the bottom and correspond to TCGA abbreviations (https://stephenturner.github.io/tcga-codes/).

STExplorer integrates GWPCA with GSEA, assigning functional insights to GWPCA results and providing spatially organised functional operations. For example, at the heterogeneous central area of section H1_4 (GG2 surrounded by benign epithelium) (Fig. 3A), GWPCA isolates LTF as the leading gene (Fig. 3A). At the same time, GSEA indicates that several immune and innate defence responses are enriched as associated functions for the same spots (Fig. 3C). Interestingly, LTF was recently shown to regulate the immune microenvironment of prostate cancer[44]. With regards to peripheral stroma that is distally located to cancerous spots within the same section, GSEA reveals functional enrichment in smooth muscle contraction, an expected observation for a non- transformed fibromuscular stroma that typically surrounds the prostate gland (Fig. 3A). Importantly, GWPCA primarily (but not exclusively) highlights the smooth muscle cell marker TAGLN2[59] as the leading gene for these peripheral stroma areas, in alignment with GSEA results. Interestingly, GWPCA also identifies TAGLN2 as a leading gene in some GG2 cancerous spots with adjacent inflamed stroma. TAGLN2 is known for its role in immune responses[60] and immune regulation in various cancers[61], supporting these findings. Finally, GWPCA assigns FABP5 and PPFIA2 as leading genes in GG4 Cribriform spots. PPFIA2 has been included in a panel of candidate genes for early detection of prostate cancer[62], while FABP5 is involved in multiple metabolic adaptations that promote prostate cancer[63, 64]. These observations represent an example of how GWPCA, together with GSEA, can be used to decipher complex spatial mosaics that associate with multifunctional leading gene expression across histological barriers.

### GWPCA leading gene expression in prostate scRNA-seq

GWPCA analyses spatially organised transcriptional variability, while single-cell approaches independently deconvolute the same variability. Due to GWPCA’s localised nature, we tested GWPCA-derived leading genes in re-analysed scRNA-seq data from the prostate cell atlas[65]. For instance in section H1_4, GWPCA leaders like FABP5 (GG4_Cribriform, Fig. 3A), extended spatial observations to the single-cell level. FABP5 was primarily expressed in luminal epithelial KLK4+ (LE-KLK4+) prostate cancer (PC) cells, linked to high magnetic resonance imaging (MRI) scores (Fig. 3D), and also in T- cells, supporting its role as an immune-metabolic marker in the tumour microenvironment (TME) that aids T cell survival through fatty acid uptake[66]. LTF (GG2 and surrounding benign epithelium leader, Fig. 3A) was mainly expressed in the club cell (CE) subpopulation (Fig. 3D), consistent with previous reports[67]. CE cells likely act as an interface between the prostate and the immune system[68], which explains the inflammation in these areas observed through functional clustering analysis (Fig. 3C). Lastly, TAGLN2 (smooth muscle leader of peripheral and tumour-surrounding inflamed stroma, Fig. 3A) was primarily expressed in fibroblasts (Fig. 3D), confirming reports of its presence in the TME, marking macrophages and cancer-associated fibroblasts[61].

Other GWPCA leading genes included KLK4 and NPY from GG2 cancerous spots of H2_1 and MPC2 and SPON2 from stroma areas of H2_5 (Sup. Fig. 5B). The oncogenic role of KLK4 is well-known and evident in luminal epithelial (LE)-KLK3^+^/KLK4^+^ PCs. With a similar expression pattern, we found NPY, a pleiotropic peptide implicated in the paracrine regulation of PC[69, 70] and MPC2, which promotes PC proliferation and growth via metabolic rewiring[71], and differentiation towards the luminal lineage[72]. Finally, SPON2 was expressed in LE-KLK3^+^ and natural killer (NK) cells, and was previously reported to promote the infiltration of the M2-polarised tumour-associated macrophages (TAMs) [73, 74]. Additionally, a positive correlation between Gleason score and average expression and/or frequency for the majority of the representative GWPCA leading genes derived from cancerous spots (Sup. Fig. 5C), overall aligns with the observed higher levels of expression in the cancerous subpopulation of cells for most patients (Sup. Fig. 5D).

These representative observations complement GWPCA/GSEA findings on the spatial data, providing independent support and additional insights at the single cell level as well as a route to combining GWPCA/GSEA with scRNA-seq analysis results.

### Diagnostic and prognostic potency of FGWC and GWPCA derived meta-signatures in prostate bulk RNA-seq

We further hypothesised that these gene leaders could form transcriptional panels with broader biological and clinical functions. We tested this by combining GWPCA-derived and NMF-derived leading gene panels into 96 meta-signatures, representing prioritised gene assemblies from cancerous, benign, or stromal tissue sections. We then analysed these meta-signatures and Seurat-derived control signatures using bulk transcriptome data and clinical information from the TCGA and GTEx consortia (see Supp. Methods).

Transcriptome analysis confirmed that most of our meta-signatures were differentially expressed in TCGA prostate tumours (PRAD) compared to matched paracancerous or normal prostate biopsies from GTEx. For instance, ten out of eleven cancerous signatures from section H1_4 were upregulated in prostate tumours with an average *log2* fold change of 1.5, while eight out of nine benign signatures were downregulated with an average *log2* fold change of -1.7 (Fig. 3E). Similar to the NMF signature results, these fold changes were consistent, indicating robust performance despite inter-patient variability. In contrast, Seurat-derived signatures showed more heterogeneous performance (Suppl. Table 7). No consistent differences in terms of performance were observed between NMF and GWPCA-derived panels or between signatures from different GWP components despite their unique gene compositions (Suppl. Fig. 6A). Similar findings were noted for all prostate sections, regardless of histological heterogeneity (Suppl. Fig. 7A), with disease and KEGG/GO analysis showing enrichment in cancer-related processes (Suppl. Fig. 6B, Suppl. Table 8). These results align with spatial GSEA findings, supporting the functional relevance of our meta-signature panels in prostate tumours.

Differential tumour expression frequently aligns with diagnostic potential. Leveraging clinical data from the TCGA PRAD panel, we extended ROC analysis to all our meta-signatures to compare their diagnostic performance. Both NMF (Fig. 3F) and GWPCA- derived (Suppl. Fig. 6C) meta-signatures showed high diagnostic power for prostate tumours, with AUC values exceeding 0.8 for cancerous gene signatures and below 0.2 (indicative of high performance for normal prostate epithelium detection) for benign gene signatures. Additionally, the deregulated expression of our meta-signatures remained statistically significant regardless of tumour stage, Gleason score, or lymph node invasion status, underscoring their suitability for early tumour diagnosis (Fig. 3G, Suppl. Fig. 6D).

Extending bulk RNA-seq analysis beyond prostate cancer, we compared differential meta-signature expression across nineteen cancer types, organised into five anatomical primary sites (Fig. 3H, Suppl. Fig. 7B, Suppl. Table 9). Unexpectedly, most benign meta-signatures showed strong downregulation in reproductive, oesophageal, and rectal neoplasms, but consistent upregulation in colonic and pancreatic tumours. Cancerous meta-signatures, however, exhibited independent clusters of overexpressed genes across all tumour types. This multi-cancer analysis highlights extensive differential expression across various primary sites, suggesting diagnostic potential beyond prostate cancer. In conclusion, STExplorer provides two complementary analytical strategies that identify differentially expressed genes with strong diagnostic and prognostic potential in bulk RNA-seq data, expanding the clinical applications of spatial transcriptome analysis.

### FGWC and GWPCA evaluate hepatic zonal patterns and disease signatures in steatotic liver disease

Steatotic liver disease (SLD) is the overarching term that describes conditions characterised by increased fat accumulation in the liver[75]. The disease spectrum ranges from isolated hepatic fat accumulation (“simple steatosis”) to a more inflammatory form (steatohepatitis) characterised by increasing scarring (hepatic fibrosis) and ultimately progression to cirrhosis, liver cancer and end-stage liver disease [76]. Key risk factors include presence of the metabolic syndrome (obesity, type 2 diabetes, high cholesterol, and hypertension), a type of SLD termed Metabolic-dysfunction Associated Steatotic Liver Disease (MASLD), and/or high alcohol intake. However, the cellular origins of the disease and the mechanisms that drive inter-patient variation in severity are not fully understood. Here, we used human liver samples from the Liver Cell Atlas[77] to demonstrate the application of the combined FGWC/GWPCA approach in a chronic disease setting. Two slices were selected: a biopsy with mild pericellular and periportal fibrosis and no steatosis (biopsy ID: JBO018), and one with 70% steatosis and clear pericellular fibrosis (biopsy ID: JBO019), referred to as “mild fibrotic” and “steatotic” respectively (see Supp. Methods).

We performed semi-supervised FGWC analysis identifying the optimum number of clusters (*k*) through STExplorer’s Fuzzy Partition Coefficient approach (Supp. Fig. 8A). Even though we used only 3 NMF factors (Supp. Fig. 8B-C), STExplorer identified five fuzzy clusters corresponding to functional units of the liver – the hepatic acini (Fig. 4A). The clusters represent hepatocytes and other cell types found in various functional zones of the liver, divided based on oxygen supply. Cluster 1 represents the central region, cluster 2 – perivenular/zone 3, cluster 3 – mid-zone/zone 2, and clusters 4 and 5 – periportal and portal regions/zone 1. The accuracy of this annotation was confirmed by the expression of known marker genes for cell types found in different zones (Supp. Fig. 8D). The clustering of spots based on the gene expression recapitulated the zonal architecture reported by Guilliams et al., 2022[77], supporting the robustness of STExplorer annotations. In both analysed samples, an additional region was identified by STExplorer, corresponding to the perivenular region (cluster 2), providing further granularity within the data absent from the original annotation. These areas consist of spots annotated as mid (mild-fibrotic sample) or as a mixture of mid/central (steatotic sample) (Fig. 4B). This presents an excellent example of the fuzzy approach in action and the idea of ecotones. The perivenular area, situated between the terminal hepatic venule and mid regions, is identified by STExplorer as an ecotone. This area includes spots where membership percentages peak, marking them mainly as pericentral (Fig. 4C-D, Supp. Fig. 8E).

**Figure 4.**
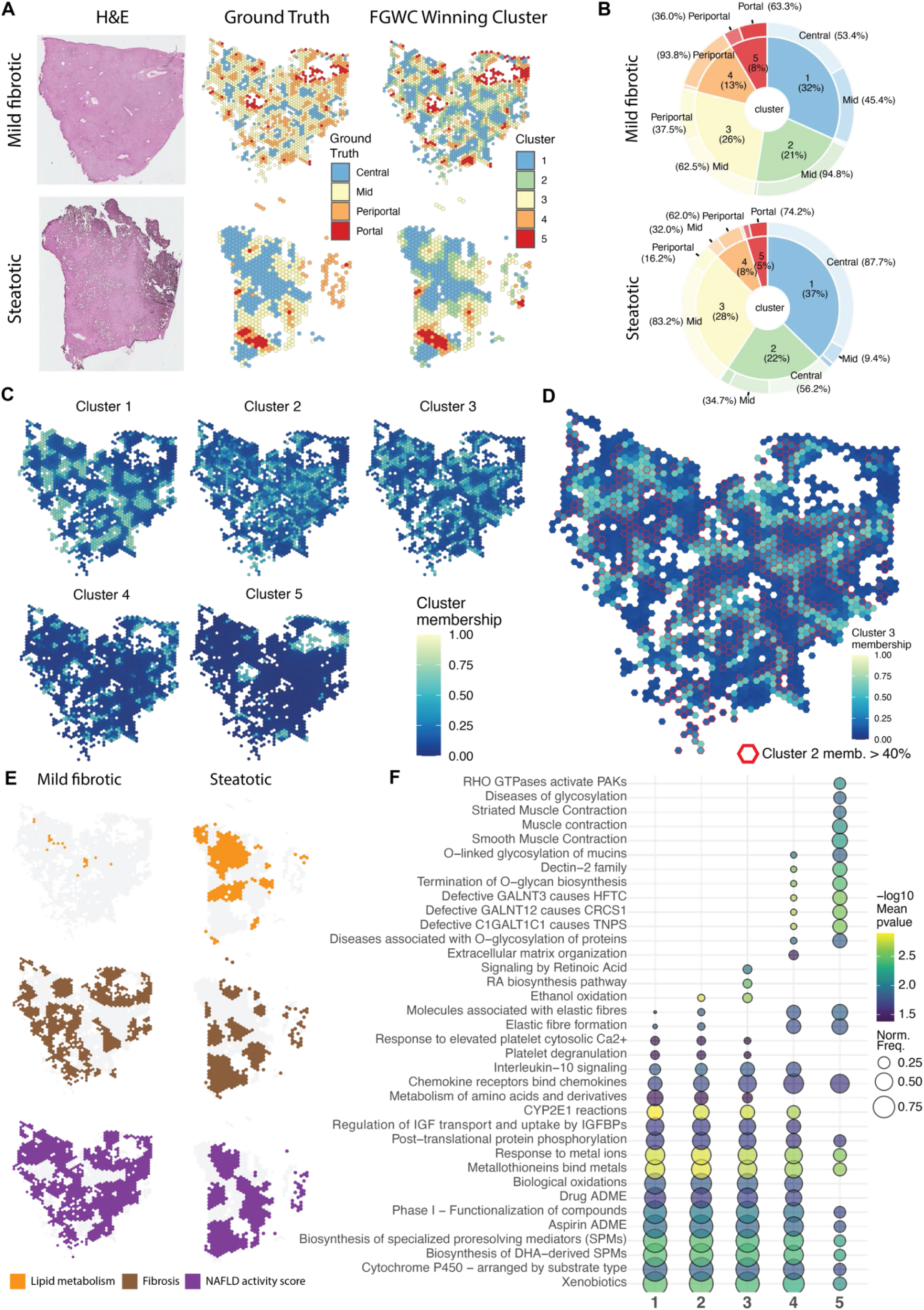
Combinatory analysis of steatotic liver with FGWC and GWPCA. A. Winning clusters from the application of FGWC in a semi-automatic way alongside the ground truth as presented in the Liver Cell Atlas and the HCE image. The colours match the ground truth to the equivalent cluster from FGWC. FGWC identified an intermediate cluster (cluster 2) as an intermediate between central and mid liver regions. **B.** Pie-doughnut charts quantitatively showcasing the match between the ground truth and FGWC clusters. The fuzziness introduced by FGWC allows for a more versatile set of clusters capturing local biology. **C.** Membership percentage maps of the Mild-fibrotic sample showcasing putative ecotones between clusters 2 and 3. **D.** Overlay of the cluster 3 membership percentages with spots of cluster 2 membership percentage > 40%. This approach helps visualise the existence of ecotones as spots with approximately mid-level percentages between the two clusters. **E.** Functional clustering of areas based on their enrichment of specific pathways using the GWPCA top leading genes in each spot. **F.** Subset (find the complete table in the supplementary figure 7) of the Reactome pathway enrichment of the Mild-fibrotic sample using the top 10 leading genes from the first three principal components (30 genes in total) in each spot.

GWPCA functional clustering of the steatotic liver sample revealed regions with enrichment in genes related to the metabolism of lipids, located primarily in acinar zones 3 and 2. The mild fibrotic sample, as expected, showed no evidence of lipid metabolism enrichment since this is a steatotic liver process (Fig. 4E, Supp. Fig. 8F). Both liver samples showed enrichment in the fibrosis[78] and in the Metabolic-dysfunction Associated Steatotic Liver Disease (MASLD) activity gene signatures[78], which include steatosis, inflammation and hepatocyte ballooning (Fig. 4E, Supp. Fig. 8F). These findings, consistent with metadata provided by Govaere *et al.*, 2020[78], further validate the ability of STExplorer to capture biological processes.

Additionally, Reactome pathway enrichment analysis using the top 10 leading genes from the first three GWPCA principal components revealed distinct pathway enrichment patterns across the five identified clusters, aligning well with liver function zonation and gradients (Fig. 4F, Supp. Fig. 8F). In the central (cluster 1) region, pathways such as xenobiotics, drug ADME (Absorption/Distribution/Metabolism/Excretion), DHA-derived SPMs (Docosahexaenoic Acid-derived Specialized Pro-Resolving Mediators), and biological oxidations are highly enriched, reflecting the central region’s role in drug metabolism[79, 80], detoxification[81], and inflammation resolution[82]. The perivenular (cluster 2) region shows high enrichment of Metallothionein-related and metal ion-related pathways, suggesting protection against metal ion toxicity alongside the expected xenobiotics pathway. The mid (cluster 3) region shows moderate enrichment of phase I - functionalisation of compounds and metal ion-related pathways, indicating an adaptive response to redox and metal ion fluctuations, supported by genes like MT1F/1G/1H/1M/1X. The periportal (cluster 4) region features xenobiotics, phase I - functionalisation of compounds, and chemokine-related pathways, with CYP2E1, CYP3A4, and CYP2A6 genes explaining roles in drug metabolism and detoxification[83–85] with chemokines indicating a putative effect of inflammatory activity in the adjacent portal areas. The portal (cluster 5) region shows pathways related to chemokine binding and smooth muscle fibre, with genes like CCL19/21, FBLN1, MFAP4, and ACTA2 indicating roles in immune cell migration[86], inflammation[87], extracellular matrix organisation[88] and fibrosis[89, 90]. The normalised pathway frequencies support the hypothesis that FGWC can substitute for ground truth when difficult to obtain, with GWPCA providing region-specific liver function insights (Fig. 4F, Supp. Fig. 8F).

### Spatially Resolved Regression Analysis Reveals the Interplay Between Senescence and Fibrosis in Pulmonary Fibrosis

Idiopathic Pulmonary Fibrosis (IPF) involves repetitive alveolar epithelial injury, driving the profibrotic reprogramming of alveolar epithelial type II (AT-II) cells into myofibroblasts via epithelial-to-mesenchymal transition (EMT)[91–93]. Myofibroblasts promote fibrosis by secreting profibrotic factors and producing excess extracellular matrix (ECM) components[94]. Additionally, during fibrosis progression, some myofibroblasts and AT- II cells acquire a senescent identity, characterised by permanent cell-cycle arrest associated with a reduction in specific cellular functions and a profibrotic secretome[95] Of interest, a recent study showed that elimination of senescent myofibroblasts with a senolytic agent alleviated bleomycin-induced pulmonary fibrosis and promoted the resolution of established fibrosis[96]. Here we utilised STExplorer’s geographically weighted regression (GWR) and spatial autocorrelation (SA) methods to further characterise the intrinsic relationship between fibrosis and senescence. GWR allows us to analyse the stationarity of a process over space by applying geographical weights before regressing locally in each neighbourhood. Our analysis of the lung sections showed that an increase in the fibrotic score (FS), which corresponds to increasing histological fibrosis severity, is accompanied by an increase in senescence (SenMayo), EMT, and ECM scores (see Supplementary Methods) (Fig. 5A, B) compared to non-fibrotic tissue (FS0; FS range: non-fibrotic (FS0), ‘mild’ (FS1), ‘moderate’ (FS2), and ‘severe’ (FS3) fibrosis)[97]. On the other hand, a TGFβ-regulated gene score, reflecting the transcriptional activity of the master regulator of fibrosis TGFβ[98], was elevated in areas of ‘mild’ fibrosis (FS1) but declined in areas with ‘moderate-severe’ fibrosis (FS2-FS3), suggesting its role in early fibrotic stages (Fig. 5A, B). Then, we utilised the Getis-Ord Gi local SA statistic to locate hot/cold spots of SenMayo score on the samples (Fig. 5C). SA tests showed that as fibrosis progresses, senescence exhibits positive SA pattern of high SenMayo scores in concentrated areas throughout the tissue (Fig. 5C) hinting towards the non-stationary nature of this process making it a good example for applying GWR.

**Figure 5.**
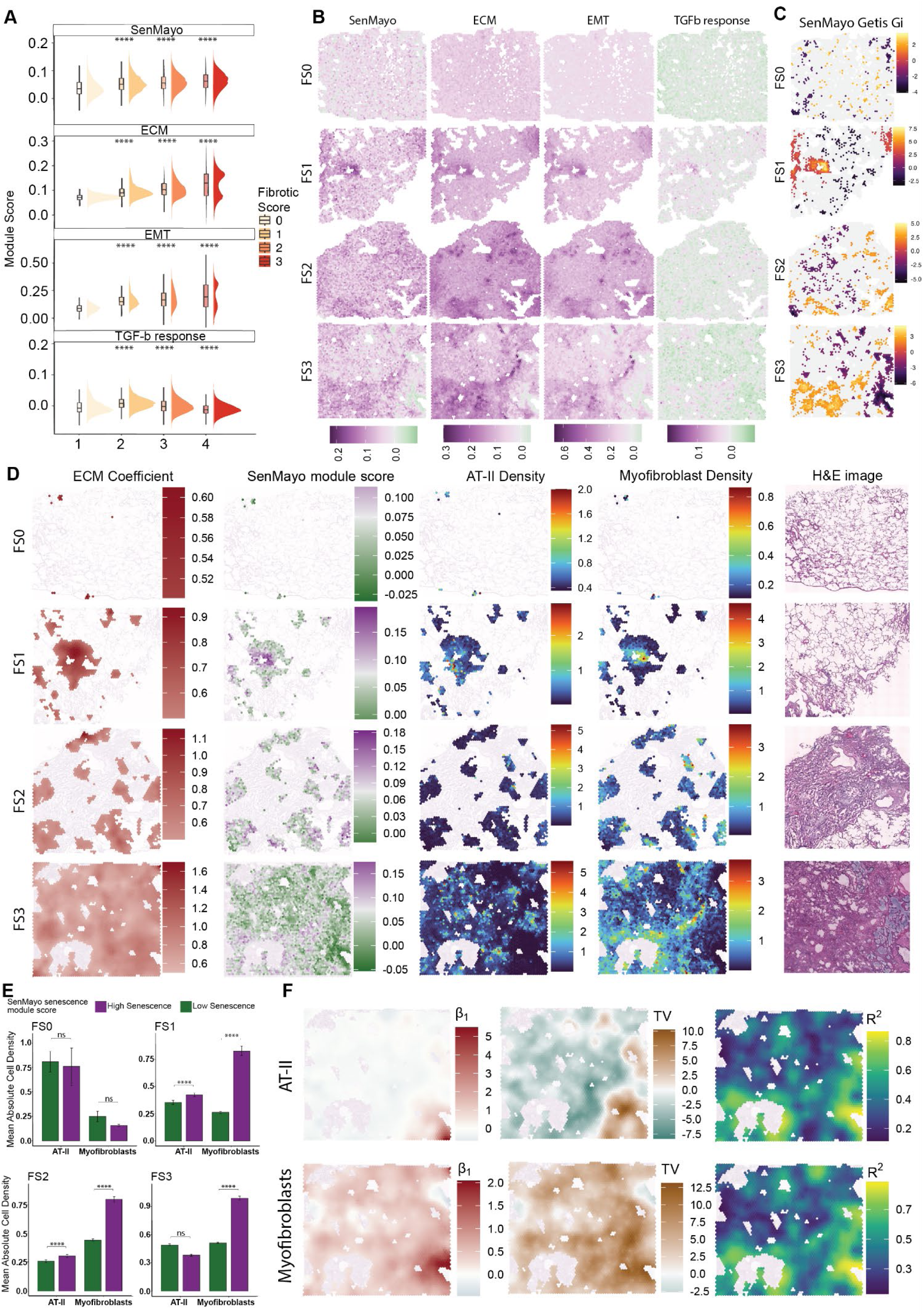
GWR analysis of spatial transcriptome data in fibrotic lung sections. A. Module scores for ECM, EMT, TGFβ-response, and SenMayo scores from all 16 samples used in the study (see Supplementary Methods). Each box plot consists of the module score values in every spot of a sample. **B.** Maps of the module scores in 4 representative samples (one from each fibrotic stage). **C.** Hot-and-Cold maps of the Getis & Ord Gi local SA statistic for the SenMayo senescence module score. The local Getis-Ord Gi captures only positive SA meaning aggregation of similar values. High positive values indicate aggregation of high senescence score values while the negative values indicate aggregation of low senescence score values. **D.** GWR results for the SenMayo∼ECM (fibrosis) local regression model. In the panel, each row is a different stage in fibrosis. Each column is composed of maps of (left to right) the ECM coefficient (filtered for areas where the coefficient was > 0.5), the SenMayo module score in areas with ECM coef. > 0.5 and coloured green to pink with pink indicating areas with high senescence, the absolute cell-type densities for AT-II and Myofibroblasts, and finally the HCE image of the sample. The cut-off for identifying the areas of high senescence was set to the 75th percentile (approx. SenMayo module score = 0.075) considering all areas of all 16 samples (see Methods). Mean absolute cell-type density in areas with ECM coef. > 0.5. Comparisons were made between areas of high and low senescence score using the same cut-off as above. Error bars show the standard error of the mean. **F.** GWR results for SenMayo∼AT-II density and SenMayo∼Myofibroblasts density regression models on the FS3 sample. The maps show results only for the areas where the SenMayo∼ECM model’s ECM coefficient (β1) is > 0.5. Top to bottom, the first row shows the coefficient values (β1) of the explanatory variables (AT-II/Myofibroblasts), the t-values (TV) of the model indicating significance, and the R-squared values. A t-value is a statistical measure that tests the significance of the local parameter estimate (coefficient) for a specific explanatory variable at each location. T-values > 1.96 or < -1.96 are considered statistically significant. ****: p <0.0001, ns: non-significant. ECM: Extra-Cellular Matrix, SenMayo: the SenMayo senescence set of genes, EMT: Epithelial to Mesenchymal Transition, TGFβ-response: genes responding to TGFb induction (more details about the scores can be found in the Methods). FS0-3: Fibrotic Score 0-3.

Using GWR, we modelled the relationship between the SenMayo and ECM scores over space. We identified senescent areas with a high positive coefficient for ECM, suggesting a parallel regulation of senescence and expression of ECM components (Fig. 5D). We observed an expansion of these areas with increasing fibrosis severity (FS1-to FS3) (Fig. 5D), implying the presence of increased numbers of senescent cells in fibrotic tissue as fibrosis progresses. To examine the cellular composition of these senescent-fibrotic areas, we determined the density of AT-II cells and myofibroblasts, two major cell populations involved in the pathogenesis of IPF. Myofibroblasts were enriched through all fibrosis stages (FS1-FS3) and exhibited higher density in areas with high SenMayo scores (Fig. 5D, E). GWR analysis modelling the SenMayo score versus AT-II/Myofibroblast density showed areas where the AT-II variable in the model had negative coefficients (i.e. when the SenMayo score increases, AT-II density is predicted to decrease). In contrast, the myofibroblast variable in these areas had positive coefficients, indicating not only the presence of myofibroblasts in senescent regions but also their opposite effect in the regression model compared to the AT-IIs (Fig. 5F). STExplorer provides the user with the means to further examine the model output. One example is the t-values (Fig. 5F) – the high absolute t-values are indicative of areas where the relationship between an explanatory variable and the dependent variable is strongly and significantly negative/positive. These localised effects, through providing insights into spatial patterns and helping identify clusters or regions of strong negative/positive influence, are crucial for understanding spatial heterogeneity and tailoring location-specific strategies.

The resolution of the 10X Visium platform, where information from each spot corresponds to multiple cells, means we cannot determine whether the cells co- expressing Senescence-Associated Secretory Phenotype (SASP) factors and ECM are myofibroblasts *per se* or neighbouring cells that may interact with myofibroblasts to promote their activation and ECM production. Future studies utilising *in situ* hybridisation and transcriptome analysis at the cellular or subcellular level are needed. Nonetheless, with STExplorer, we demonstrated that GWR and SA methods provide critical insights into the spatial dynamics of senescence and fibrosis in IPF, revealing the evolving interplay between senescence, ECM accumulation, and cellular composition across fibrotic stages.

## Methods

### Fuzzy Geographically Weighted Clustering

The idea of FGWC was first presented in geodemographics by Feng and Flowerdew[99] as a post-analysis adjustment of group memberships. However, FGWC embeds geographic considerations into the core of the clustering process while it adjusts the membership values considering “neighbourhood effects” and ensures that geographical context is not overlooked. Additionally, by incorporating principles from geographical spatial interaction models[100], FGWC proposes a weighting mechanism that not only accounts for variable values but also considers the spatial relationships between areas, thereby proposing “geographically aware” cluster centres[27].

FGWC applies weights to membership values based on spatial interactions. This step is crucial because it makes the algorithm sensitive to both the intrinsic data characteristics and the spatial layout of the areas. The degree of fuzziness, controlled by the fuzzy exponent, further influences this process, suggesting that the integration of geographical effects into the iterative cycle will indeed reshape cluster centres to reflect their geographical context more accurately[27]. By doing so, FGWC becomes a location- sensitive spatial omics toolset.

### Internal mechanism

The first step before running FGWC is to find a lower-dimensional representation of the data using Non-negative Matrix Factorisation (NMF). With NMF, the (`genes x locations`) gene expression matrix *A*, is factorised so that two new matrices W and H are generated so that the product WH approximates *A* (Equation 1).3

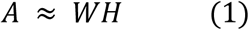

In equation (4), *W* is the (`genes x factors`) feature loading matrix, and *W* the (`factors x locations`). STExplorer uses the scater R package to calculate the NMF function which uses the C++ implementation of NMF in R from the RcppML R package[101]. Then, the reduced dimensions matrix *W* is used as input by FGWC.

The FGWC mechanism is explained in detail by Mason et. al.[27]. The model considers the effect of one region on another as the product of the population sizes of the regions. In the current STExplorer implementation, the population of each area is considered to be similar (equal to one cell per area). This interaction is moderated by a distance decay factor, which acts as the divisor. The fuzzy membership matrix is subsequently updated by incorporating the geographical weight during each iteration of clustering, as shown in the following equation:

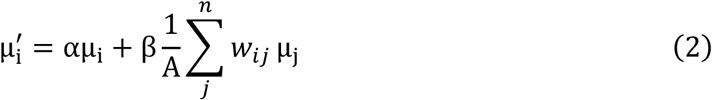

In this formulation, μ^′^ represents the updated fuzzy membership of the area i, while μ_i_ is the prior fuzzy membership of the same area. The variable *n* denotes the number of areas considered. The term *W*_*ij*_ is the interaction weighting measure between two geographic areas *i* and *j*. This weight is influenced by the distance between the centres of the areas and their populations. The parameter *A* is chosen to ensure that the average of the weighted membership values remains within the range [0, 1] [27].

The interaction weight *W*_*ij*_ is defined as:

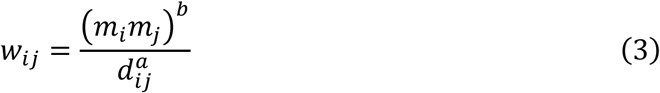

where *m*_*i*_and *m*_*i*_are the populations of areas *i* and *j*, respectively, *d*_*ij*_is the distance between the centres of areas *i* and *j*. Finally, the user-defined parameters *a* and *b* control the influence of distance and population respectively on the interaction weight[27].

The *α* and *β* from equation (2) are the weights applied to the old fuzzy membership value and the mean of membership values of neighbourhood areas respectively and are calculated as follows[27]:

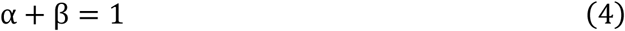

STExplorer’s implementation of FGWC is based on the *naspaclust* R package[102]. The default setup for STExplorer’s FGWC is the classic approach by Mason *et. al.*[27], as explained above with the population in each location set to be equal to one cell.

### Geographically Weighted Principal Component Analysis

Geographically Weighted Principal Component Analysis (GWPCA) is a method developed and used in Geographical sciences to study spatial effects that are often vital to a more complete understanding of a given process[26]. GWPCA extends the traditional PCA by integrating spatial variability into the analysis. This method facilitates the exploration of locale-specific variability within multidimensional datasets, adapting PCA to consider the spatial layout of data points in its computations. GWPCA incorporates spatial weighting, meaning closer neighbours have a greater influence on the analysis of each observation (location or spot). Additionally, unlike traditional PCA, which provides global principal components, GWPCA provides principal components that are locally derived. GWPCA utilises a moving window approach – starting from one location in the dataset, setting it in the centre and placing a bandwidth-defined window around it. The observations in the window are down-weighted according to their distance from the central location – decaying to close to zero for the furthest observations. A PCA is completed based on these weighted observations. Once the calculations are completed for this location, the window moves to the next and the process is repeated until all locations have been visited. As a result, each location has its own set of principal components that reflect the local variation in the data. A spatial weighting scheme (e.g., Gaussian kernel) is used to determine the influence of neighbouring points on the analysis of a particular observation. The bandwidth of the spatial kernel controls the degree of localisation.

The GWPCA output follows the below structure:

1. Local Scores: the scores of each location in its local principal components.
2. Local Loadings: the loadings of each gene in each principal component for each location
3. Local Explained Variance: proportion of variance explained by each local principal component.
4. Global Components: Global components for comparison with traditional PCA results.

The GWPCA output includes a set of local leading scores for the variables (genes) and local scores in each Principal Component (PC) for each location alongside local percentage variation metrics. Each gene gets a leading score in each principal component in each location indicating the level of its contribution in the variability explained by a specific PC in a certain locality. This information is later passed down to secondary analyses to reveal the locally affected biology and increase the biological utility of the GWPCA results. Utilising the leading scores within STExplorer, ranked lists of genes can be generated. Through this approach, we can investigate the functional impact of these leading genes locally. The ranked lists are then used to perform functional annotation per location. Subsequently, these annotations cluster together locations that behave in a similar way and identify which processes or pathways are affected in these locations. Currently, STExplorer implements Gene Set Enrichment Analysis (GSEA) using the Molecular Signatures Database (MSigDB) to achieve this. In general, though, any form of gene-related annotation can be used.

### Internal mechanism

The `gwpcaSTE` function first calculates the distance matrix `dMat` using the location coordinates and the `p` parameter to calculate the distance between locations. The p parameter defaults to 2 which results in calculating the Euclidean distance which equates to the physical distance between locations. The function then calculates the distance weights `wt` using a kernel function and the distance matrix. For a target location *s*_*i*_ and a set of neighbouring observations *s*_1_, *s*_2_, …, *s*_*n*_, we define the weight for a neighbouring location *s*_*i*_ as *W*_*i*_. This weight can be calculated using a kernel function

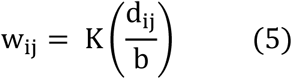

where *d*_*ij*_is the distance between *s*_*i*_and *s*_*i*_, and *b* is a bandwidth parameter that controls the spatial extent of the local analysis. Common choices for *K* include the Gaussian (6) and the Exponential (7) kernels:

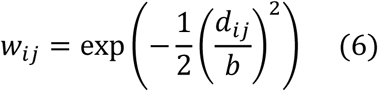

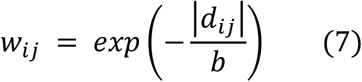

The data matrix `x` includes the gene expression for a location *s*_*i*_ and its neighbours. The weighted data matrix `x_weighted` is calculated by multiplying the data matrix `x` with the specific weights `wt` for location *s*_*i*_ and its neighbours. Then a Singular Value Decomposition (SVD) on the weighted data matrix `x_weighted` is performed which extracts the principal components from the data. The top `k` principal components are retained, and the scores for each location are calculated using the retained principal components. After completing this step for all locations, we are left with two three-dimensional arrays. One includes the leading scores of each gene, in each location, for each PC (with dimensions: `locations x genes x PCs`). The other contains the PC scores of each location for each local PCA run (with dimensions: `locations x PCs x locations`). These two arrays of eigenvalues and eigenvectors are then utilised for the downstream analyses.

### Empirical running time

In supplementary table 1 (Supp. Table 1) we provide a set of running times for different computer setups and parameters. In general, increasing the number of HVGs used in GWPCA drastically increases the time required by GWPCA to complete the task. On average, when using the top 10% HVGs (200-400 genes) for 2500 – 4000 locations, GWPCA takes between 4 and 20 minutes. Additionally, for a five-sample analysis running in parallel, approximately 100 GB of RAM is required.

### Geographically Weighted Regression

Geographically Weighted Regression (GWR)[13] is an extension of traditional regression that accounts for spatial heterogeneity, making it particularly useful for exploratory spatial data analysis. STExplorer, for now, utilises the basic GWR implementation as presented in the GWmodel R package[38]. Unlike conventional regression models, which assume that relationships between variables are uniform across the study area, GWR allows these relationships to vary locally. This flexibility facilitates a deeper understanding of spatially varying processes and relationships within complex datasets. When exploring GWR results, we usually plot the resulting local regression coefficient estimates as well as the (pseudo) t-values to provide evidence of non-stationarity.

In a standard regression model with one predictor, the relationship is defined as:

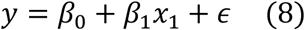

where *y* is the dependent variable, *x*_1_is the independent variable, *β*_0_and *β*_1_are the regression coefficients, and *∈* represent the error term. In GWR, the model is redefined to account for spatial variability:

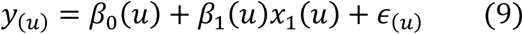

Here, *u* represents the spatial coordinates of a location, and the regression coefficients (*β*_0_(*u*), *β*_1_(*u*)) are calculated locally for each location *u*. These coefficients vary depending on spatial proximity, achieved through a weighting scheme that assigns higher weights to data points closer to the location *u*.

To implement GWR, a spatial kernel function, such as Gaussian or Exponential, is applied to calculate weights based on distances between the focal point *u* and neighbouring locations. The weights are determined as:

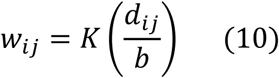

where *d*_*i*_is the distance between locations *i*, *j*, and *b* is a bandwidth parameter controlling the spatial influence, and *K* is the kernel function (e.g., Gaussian kernel). The localised regression coefficients are calculated for each location *u* by solving the weighted least squares problem using the spatially weighted data. For a deep dive into GWR, we suggest looking into Brunsdon, *et al.*,[13], Gollini, *et al.*,[38], and Comber *et al.*[103].

### Spatial Autocorrelation

True to not reinventing the wheel, STExplorer builds on the *spdep* R package[104, 105] to allow the users to identify spatially variable genes or features utilising any of the SA statistics shown here: Global Moran’s *I*[28, 106, 107]:

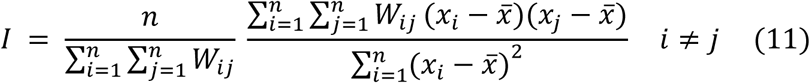

Global Geary’s *C*[29, 106, 107]:

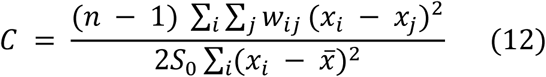

Global Getis and Ord’s *G*[30]:

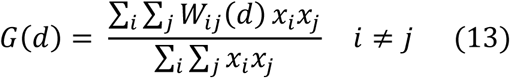

Local Moran’s *Ii*[31]:

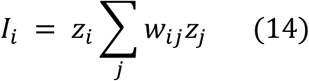

Local Geary’s *Ci*[31]:

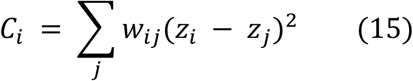

Local Getis and Ord’s *Gi*[32]:

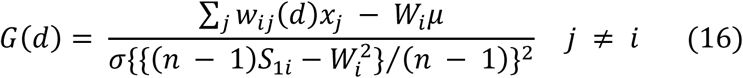

A deeper understanding of the internal mechanics of each statistical method can be found in their respective original publications. Additionally, in a previous publication[14], we attempted to show the biological significance of SA.

## Discussion

Spatial transcriptomics has transformed our ability to investigate tissue organisation and gene expression, providing unprecedented spatial context to biological phenomena. However, existing analytical frameworks often fail to exploit the spatial dimension fully, limiting the scope of insights that can be derived from spatial omics. Here, we introduce STExplorer, an R package that bridges computational geography and bioinformatics, providing a novel, spatially-aware analytical framework for exploring spatial omics data.

STExplorer leverages methods with a long history of refinement in computational geography, such as Geographically Weighted Principal Component Analysis (GWPCA), Fuzzy Geographically Weighted Clustering (FGWC), Geographically Weighted Regression (GWR), and Spatial Autocorrelation (SA) calculations. By adapting these tools to spatial transcriptomics, STExplorer enables localised, context-specific analyses that reveal spatial heterogeneity and intricate biological patterns often overlooked by global approaches. The visualisation functions included in STExplorer empower biologists to effectively interpret and communicate their findings, enabling the exploration of spatial relationships within tissue microenvironments and bridging the gap between a package’s output and its practical application.

The use of SpatialFeatureExperiment (SFE) objects ensures compatibility with existing Bioconductor pipelines and offers standardised encoding of spatial data, enhancing interoperability. This design philosophy underscores the importance of integrating spatial biology with established tools from computational geography, allowing researchers to focus on biological discovery without the need to “reinvent the wheel.”

Through case studies in tumours, chronic disease, senescence and organ development, we demonstrated STExplorer’s ability to uncover biologically relevant insights, such as spatially interpretable clusters, spatially autocorrelated features, local sources of variability, usually leading to generating marker genes or gene signatures, as well as identifying spatial heterogeneity in relationships between biological variables. These findings highlight the value of spatially informed analyses in understanding cellular interactions and tissue architecture.

However, the current implementation of STExplorer is not without limitations. While it provides robust tools for spatial analysis, its reliance on computational geography methods necessitates a basic understanding of these techniques, which could pose a learning curve for biologists new to this field. In addition, the growing size of datasets means that STExplorer’s analyses will require more time and computing resources to complete. To this end, we are committed to continuing the development and refinement of the methods to tackle these issues.

Looking forward, the development of STExplorer opens several avenues for further research. The integration of additional spatial methods, such as spatially variable gene detection and multi-modal data analysis, could broaden its utility.

In conclusion, STExplorer provides a powerful, flexible, and user-friendly platform for analysing spatial transcriptomics data, bridging the gap between computational geography and spatial biology as well as analysis output and biological application. By addressing spatial heterogeneity in biological data, STExplorer offers new opportunities to advance our understanding of tissue biology and cellular interactions in health and disease.

## Data availability

The prostate data originate from Erickson A. *et. al.*[42]. Count matrices, high-resolution histological images and additional material were downloaded from Mendeley Data (https://doi.org/10.17632/svw96g68dv.1).

The liver data were downloaded from the Liver Cell Atlas (https://livercellatlas.org/index.php).

The developmental eye data were kindly contributed to this work by Dorgau B. *et. al.*[46], and can be found in Gene Expression Omnibus (GEO) under accession number GSE234971.

The human lung fibrosis data originate from Franzén L *et al.*[97] and are accessible from the EBI BioStudies portal at https://www.ebi.ac.uk/biostudies/studies/S-BSST1409

## Code availability

STExplorer is currently released as an open-source R package on GitHub. The source code for STExplorer can be accessed, downloaded, and installed from https://github.com/LefterisZ/STExplorer. The repository also includes detailed vignettes with examples of usage. The analysis for this work is available in a separate repository, and can be accessed from https://github.com/ncl-icbam/STExplorer_Analysis.

## Acknowledgements

EZ is supported by a studentship from the Medical Research Council Discovery Medicine North (DiMeN) Doctoral Training Partnership (grant number: MR/N013840/1). STC is supported by the British Heart Foundation (PG/23/11093) and the Royal Society (RG/R1/241197). RQ & ML would like to acknowledge funding from BBSRC (BB/T004460/1). QMA is an NIHR Senior Investigator and is supported by the Newcastle NIHR Biomedical Research Centre, and the Horizon Europe, IMI2 and IHI research and innovation programmes of the European Union under Grant Agreements 777377 (LITMUS), 101132901 (LIVERAIM), 101136259 (EDC-MASLD) and 101136622 (THRIVE).

AG would like to thank the European Regional Development Fund of the European Union along with the National Operational Program Competitiveness, Entrepreneurship and Innovation (MIS 5050733) and Fondation Sante (5935) for their financial support. Views and opinions expressed are those of the authors and do not necessarily reflect those of the European Union, EFPIA, the NHS, NIHR, or the UK Department of Health.

## Author Contributions

Conceptualization: EZ and SJC; Data Curation: EZ and AG; Formal Analysis: EZ, AR, AU and AG; Funding Acquisition: SJC; Investigation: EZ, NIV, AR, AU, STC, DT, QMA, AG and SJC; Methodology: EZ, RQ, AC and SJC; Project Administration: EZ and SJC; Resources: BD, ML and DT; Software: EZ, AR, AU and AG; Supervision: RQ, AC and SJC; Validation: EZ, STC, RQ, ML, QMA, AG and SJC; Visualization: EZ, AR, AU, AG and SJC; Writing – Original Draft Preparation: EZ, NIV, AR, AG and SJC; Writing – Review and Editing: All.

## References

1. Zhuang, X., Spatially resolved single-cell genomics and transcriptomics by imaging. Nature Methods, 2021. 18(1): p. 18–22.

2. Femino, A.M., et al., Visualization of Single RNA Transcripts in Situ. Science, 1998. 280(5363): p. 585-590.

3. Xia, C., et al., Spatial transcriptome profiling by MERFISH reveals subcellular RNA compartmentalization and cell cycle-dependent gene expression. Proceedings of the National Academy of Sciences, 2019. 116(39): p. 19490–19499.

4. Wang, X., et al., Three-dimensional intact-tissue sequencing of single-cell transcriptional states. Science, 2018. 361(6400).

5. Zollinger, D.R., et al., GeoMx™ RNA Assay: High Multiplex, Digital, Spatial Analysis of RNA in FFPE Tissue, in In Situ Hybridization Protocols, B.S. Nielsen and J. Jones, Editors. 2020, Springer US: New York, NY. p. 331-345.

6. Papalexi, E. and R. Satija, Single-cell RNA sequencing to explore immune cell heterogeneity. Nature Reviews Immunology, 2018. 18(1): p. 35–45.

7. Ståhl, P.L., et al., Visualization and analysis of gene expression in tissue sections by spatial transcriptomics. Science, 2016. 353(6294): p. 78-82.

8. Rodriques, S.G., et al., Slide-seq: A scalable technology for measuring genome-wide expression at high spatial resolution. Science, 2019. 363(6434): p. 1463- 1467.

9. Stickels, R.R., et al., Highly sensitive spatial transcriptomics at near-cellular resolution with Slide-seqV2. Nat Biotechnol, 2021. 39(3): p. 313–319.

10. Chen, A., et al., Spatiotemporal transcriptomic atlas of mouse organogenesis using DNA nanoball-patterned arrays. Cell, 2022. 185(10): p. 1777–1792.e21.

11. Openshaw, S. and P.J. Taylor, A million or so correlation coefficients : three experiments on the modifiable areal unit problem, in Statistical Applications in the Spatial Sciences, N. Wrigley, Editor. 1979, Pion: London. p. 127-144.

12. Tobler, W.R., A Computer Movie Simulating Urban Growth in the Detroit Region. Economic Geography, 1970. 46(sup1): p. 234–240.

13. Brunsdon, C., A.S. Fotheringham, and M.E. Charlton, Geographically Weighted Regression: A Method for Exploring Spatial Nonstationarity. Geographical Analysis, 1996. 28(4): p. 281–298.

14. Zormpas, E., et al., Mapping the transcriptome: Realizing the full potential of spatial data analysis. Cell, 2023. 186(26): p. 5677–5689.

15. Fu, W., et al., Peri-urbanization may vary with vegetation restoration: A large scale regional analysis. Urban Forestry & Urban Greening, 2018. 29: p. 77–87.

16. Granger, S.J., et al., Phosphate stable oxygen isotope variability within a temperate agricultural soil. Geoderma, 2017. 285: p. 64–75.

17. Lu, B., et al., Uncovering drivers of community-level house price dynamics through multiscale geographically weighted regression: A case study of Wuhan, China. Spatial Statistics, 2023. 53: p. 100723.

18. Liu, Y., et al., Quantifying the spatio-temporal drivers of planned vegetation restoration on ecosystem services at a regional scale. Science of The Total Environment, 2019. 650: p. 1029–1040.

19. Fotheringham, A.S., C. Brunsdon, and M. Charlton, Quantitative Geography: Perspectives on Spatial Data Analysis. 2000: SAGE.

20. Corcoran, J., et al., The use of spatial analytical techniques to explore patterns of fire incidence: A South Wales case study. Computers, Environment and Urban Systems, 2007. 31(6): p. 623–647.

21. Grimes, M., et al., Land cover changes across Greenland dominated by a doubling of vegetation in three decades. Scientific Reports, 2024. 14(1): p. 3120.

22. Demšar, U., et al., Principal Component Analysis on Spatial Data: An Overview. Annals of the Association of American Geographers, 2013. 103(1): p. 106–128.

23. Fotheringham, A.S. and C. Brunsdon, Local Forms of Spatial Analysis. Geographical Analysis, 1999. 31(4): p. 340–358.

24. Comber, A., et al., A Route Map for Successful Applications of Geographically Weighted Regression. Geographical Analysis, 2023. 55(1): p. 155–178.

25. Comber, A.J., P. Harris, and N. Tsutsumida, Improving land cover classification using input variables derived from a geographically weighted principal components analysis. ISPRS Journal of Photogrammetry and Remote Sensing, 2016. 119: p. 347–360.

26. Harris, P., C. Brunsdon, and M. Charlton, Geographically weighted principal components analysis. International Journal of Geographical Information Science, 2011. 25(10): p. 1717–1736.

27. Mason, G. and R. Jacobson. *Fuzzy geographically weighted clustering*. in *Proceedings of the 9th International Conference on Geocomputation*. 2007. Maynooth Eire, Ireland.

28. Moran, P.A.P., Notes on Continuous Stochastic Phenomena. Biometrika, 1950. 37(1/2): p. 17–23.

29. Geary, R.C., The Contiguity Ratio and Statistical Mapping. The Incorporated Statistician, 1954. 5(3): p. 115–146.

30. Getis, A. and J.K. Ord, The Analysis of Spatial Association by Use of Distance Statistics. Geographical Analysis, 1992. 24(3): p. 189–206.

31. Anselin, L., Local Indicators of Spatial Association—LISA. Geographical Analysis, 1995. 27(2): p. 93–115.

32. Ord, J.K. and A. Getis, Local Spatial Autocorrelation Statistics: Distributional Issues and an Application. Geographical Analysis, 1995. 27(4): p. 286–306.

33. Lambda, M., H. Alik, and P. Lior, *SpatialFeatureExperiment: Integrating SpatialExperiment with Simple Features in sf*. 2022.

34. Pebesma, E., Simple Features for R: Standardized Support for Spatial Vector Data. R Journal, 2018. 10(1): p. 439–446.

35. McCarthy, D.J., et al., Scater: pre-processing, quality control, normalization and visualization of single-cell RNA-seq data in R. Bioinformatics, 2017. 33(8): p. 1179–1186.

36. Hao, Y., et al., Dictionary learning for integrative, multimodal and scalable single- cell analysis. Nature Biotechnology, 2024. 42(2): p. 293–304.

37. Lun, A.T., D.J. McCarthy, and J.C. Marioni, A step-by-step workflow for low-level analysis of single-cell RNA-seq data with Bioconductor. F1000Res, 2016. 5: p. 2122.

38. Gollini, I., et al., GWmodel: An R Package for Exploring Spatial Heterogeneity Using Geographically Weighted Models. Journal of statistical software, 2015. 63.

39. Stan, O., *Developing GIS-relevant zone-based spatial analysis methods*. Spatial analysis: modelling in a GIS environment, 1996: p. 55-73.

40. Bezdek, J.C., R. Ehrlich, and W. Full, FCM: The fuzzy c-means clustering algorithm. Computers & Geosciences, 1984. 10(2): p. 191–203.

41. Arnot, C., et al., Landscape metrics with ecotones: pattern under uncertainty. Landscape Ecology, 2004. 19(2): p. 181–195.

42. Erickson, A., et al., Spatially resolved clonal copy number alterations in benign and malignant tissue. Nature, 2022. 608(7922): p. 360-367.

43. Kim, H. and H. Park, Sparse non-negative matrix factorizations via alternating non- negativity-constrained least squares for microarray data analysis. Bioinformatics, 2007. 23(12): p. 1495–1502.

44. Zhao, Q., Y. Cheng, and Y. Xiong, *LTF Regulates the Immune Microenvironment of Prostate Cancer Through JAK/STAT3 Pathway*. Front Oncol, 2021. 11: p. 692117.

45. Riedel, M., et al., In vivo CRISPR inactivation of Fos promotes prostate cancer progression by altering the associated AP-1 subunit Jun. Oncogene, 2021. 40(13): p. 2437-2447.

46. Dorgau, B., et al., Single-cell analyses reveal transient retinal progenitor cells in the ciliary margin of developing human retina. Nature Communications, 2024. 15(1): p. 3567.

47. Cheng, B., et al., MT1G, an emerging ferroptosis-related gene: A novel prognostic biomarker and indicator of immunotherapy sensitivity in prostate cancer. Environ Toxicol, 2024. 39(2): p. 927-941.

48. Henrique, R., et al., MT1G hypermethylation is associated with higher tumor stage in prostate cancer. Cancer Epidemiol Biomarkers Prev, 2005. 14(5): p. 1274–8.

49. Bonaccorsi, L., et al., Androgen receptor regulation of the seladin-1/DHCR24 gene: altered expression in prostate cancer. Lab Invest, 2008. 88(10): p. 1049-56.

50. Carbonetti, G., et al., FABP5 coordinates lipid signaling that promotes prostate cancer metastasis. Sci Rep, 2019. 9(1): p. 18944.

51. Langlieb, J., et al., The cell type composition of the adult mouse brain revealed by single cell and spatial genomics. bioRxiv, 2023: p. 2023.03.06.531307.

52. Sagredo, A.I., et al., TRPM4 regulates Akt/GSK3-β activity and enhances β-catenin signaling and cell proliferation in prostate cancer cells. Mol Oncol, 2018. 12(2): p. 151-165.

53. Berg, K.D., et al., TRPM4 protein expression in prostate cancer: a novel tissue biomarker associated with risk of biochemical recurrence following radical prostatectomy. Virchows Arch, 2016. 468(3): p. 345–55.

54. Schröder, S.K., M. Pinoé-Schmidt, and R. Weiskirchen, Lipocalin-2 (LCN2) Deficiency Leads to Cellular Changes in Highly Metastatic Human Prostate Cancer Cell Line PC-3. Cells, 2022. 11(2).

55. Tung, M.C., et al., Knockdown of lipocalin-2 suppresses the growth and invasion of prostate cancer cells. Prostate, 2013. 73(12): p. 1281–90.

56. Barer, L., et al., Lipocalin-2 regulates the expression of interferon-stimulated genes and the susceptibility of prostate cancer cells to oncolytic virus infection. Eur J Cell Biol, 2023. 102(2): p. 151328.

57. Kogan-Sakin, I., et al., Prostate stromal cells produce CXCL-1, CXCL-2, CXCL-3 and IL-8 in response to epithelia-secreted IL-1. Carcinogenesis, 2009. 30(4): p. 698-705.

58. Guo, C., et al., Targeting myeloid chemotaxis to reverse prostate cancer therapy resistance. Nature, 2023. 623(7989): p. 1053-1061.

59. Tsuji-Tamura, K., et al., The canonical smooth muscle cell marker TAGLN is present in endothelial cells and is involved in angiogenesis. Journal of Cell Science, 2021. 134(15).

60. Kim, H.-R., et al., Transgelin-2: A Double-Edged Sword in Immunity and Cancer Metastasis. Frontiers in Cell and Developmental Biology, 2021. 9.

61. Pan, T., S. Wang, and Z. Wang, An Integrated Analysis Identified TAGLN2 As an Oncogene Indicator Related to Prognosis and Immunity in Pan-Cancer. Journal of Cancer, 2023. 14(10): p. 1809–1836.

62. Leyten, G.H.J.M., et al., Identification of a Candidate Gene Panel for the Early Diagnosis of Prostate Cancer. Clinical Cancer Research, 2015. 21(13): p. 3061–3070.

63. Pujana-Vaquerizo, M., L. Bozal-Basterra, and A. Carracedo, Metabolic adaptations in prostate cancer. British Journal of Cancer, 2024. 131(8): p. 1250–1262.

64. O’Sullivan, S.E. and M. Kaczocha, FABP5 as a novel molecular target in prostate cancer. Drug Discovery Today, 2020. 25(11): p. 2056–2061.

65. Tuong, Z.K., et al., Resolving the immune landscape of human prostate at a single- cell level in health and cancer. Cell Rep, 2021. 37(12): p. 110132.

66. Pan, Y., et al., Survival of tissue-resident memory T cells requires exogenous lipid uptake and metabolism. Nature, 2017. 543(7644): p. 252-256.

67. Song, H., et al., Single-cell analysis of human primary prostate cancer reveals the heterogeneity of tumor-associated epithelial cell states. Nat Commun, 2022. 13(1): p. 141.

68. Kiviaho, A., et al., Immunosuppression in the prostate tumor microenvironment is tied to androgen deprivation therapy-resistant club-like cells. bioRxiv, 2024: p. 2024.03.25.586330.

69. Ding, Y., et al., Neuropeptide Y nerve paracrine regulation of prostate cancer oncogenesis and therapy resistance. The Prostate, 2021. 81(1): p. 58–71.

70. Sigorski, D., et al., Neuropeptide Y and its receptors in prostate cancer: associations with cancer invasiveness and perineural spread. Journal of Cancer Research and Clinical Oncology, 2023. 149(9): p. 5803–5822.

71. Bader, D.A., et al., Mitochondrial pyruvate import is a metabolic vulnerability in androgen receptor-driven prostate cancer. Nature Metabolism, 2019. 1(1): p. 70–85.

72. Giafaglione, J.M., et al., Prostate lineage-specific metabolism governs luminal differentiation and response to antiandrogen treatment. Nature Cell Biology, 2023. 25(12): p. 1821–1832.

73. Huang, C., et al., Tumor cell-derived SPON2 promotes M2-polarized tumor- associated macrophage infiltration and cancer progression by activating PYK2 in CRC. Journal of Experimental & Clinical Cancer Research, 2021. 40(1): p. 304.

74. Lucarelli, G., et al., Spondin-2, a Secreted Extracellular Matrix Protein, is a Novel Diagnostic Biomarker for Prostate Cancer. J. Urol., 2013.

75. Rinella, M.E., et al., A multisociety Delphi consensus statement on new fatty liver disease nomenclature. J Hepatol, 2023. 79(6): p. 1542–1556.

76. Anstee, Q.M., et al., From NASH to HCC: current concepts and future challenges. Nat Rev Gastroenterol Hepatol, 2019. 16(7): p. 411–428.

77. Guilliams, M., et al., Spatial proteogenomics reveals distinct and evolutionarily conserved hepatic macrophage niches. Cell, 2022. 185(2): p. 379–396.e38.

78. Govaere, O., et al., Transcriptomic profiling across the nonalcoholic fatty liver disease spectrum reveals gene signatures for steatohepatitis and fibrosis. Sci Transl Med, 2020. 12(572).

79. Sturgill, M.G. and G.H. Lambert, Xenobiotic-induced hepatotoxicity: mechanisms of liver injury and methods of monitoring hepatic function. Clinical Chemistry, 1997. 43(8): p. 1512–1526.

80. Talevi, A. and C.L. Bellera, Drug Metabolism Functionalization (Phase I) Reactions, in The ADME Encyclopedia: A Comprehensive Guide on Biopharmacy and Pharmacokinetics. 2021, Springer International Publishing: Cham. p. 1-7.

81. Conde de la Rosa, L., et al., Role of Oxidative Stress in Liver Disorders. Livers, 2022. 2(4): p. 283–314.

82. Han, Y.H., et al., Specialized Proresolving Mediators for Therapeutic Interventions Targeting Metabolic and Inflammatory Disorders. Biomol Ther (Seoul), 2021. 29(5): p. 455–464.

83. Ma, R., et al., Metabolic and non-metabolic liver zonation is established non- synchronously and requires sinusoidal Wnts. eLife, 2020. 9: p. e46206.

84. Albadry, M., et al., Cross-species variability in lobular geometry and cytochrome P450 hepatic zonation: insights into CYP1A2, CYP2D6, CYP2E1 and CYP3A4. Frontiers in Pharmacology, 2024. **15**.

85. Panday, R., C.P. Monckton, and S.R. Khetani, The Role of Liver Zonation in Physiology, Regeneration, and Disease. Semin Liver Dis, 2022. 42(1): p. 1–16.

86. Yan, Y., et al., CCL19 and CCR7 Expression, Signaling Pathways, and Adjuvant Functions in Viral Infection and Prevention. Frontiers in Cell and Developmental Biology, 2019. 7.

87. Kozai, M., et al., Essential role of CCL21 in establishment of central self-tolerance in T cells. Journal of Experimental Medicine, 2017. 214(7): p. 1925–1935.

88. Uzzaman, A., et al., Discovery of small extracellular vesicle proteins from human serum for liver cirrhosis and liver cancer. Biochimie, 2020. 177: p. 132–141.

89. Iakovleva, V., et al., Mfap4: a promising target for enhanced liver regeneration and chronic liver disease treatment. npj Regenerative Medicine, 2023. 8(1): p. 63.

90. Rockey, D.C., N. Weymouth, and Z. Shi, Smooth Muscle α Actin (Acta2) and Myofibroblast Function during Hepatic Wound Healing. PLOS ONE, 2013. 8(10): p. e77166.

91. Moss, B.J., S.W. Ryter, and I.O. Rosas, Pathogenic Mechanisms Underlying Idiopathic Pulmonary Fibrosis. Annu Rev Pathol, 2022. 17: p. 515–546.

92. Martinez, F.J., et al., Idiopathic pulmonary fibrosis. Nat Rev Dis Primers, 2017. 3: p. 17074.

93. Kim, K.K., et al., Alveolar epithelial cell mesenchymal transition develops in vivo during pulmonary fibrosis and is regulated by the extracellular matrix. Proc Natl Acad Sci U S A, 2006. 103(35): p. 13180–5.

94. Younesi, F.S., et al., Fibroblast and myofibroblast activation in normal tissue repair and fibrosis. Nat Rev Mol Cell Biol, 2024. 25(8): p. 617–638.

95. Schafer, M.J., et al., Cellular senescence mediates fibrotic pulmonary disease. Nat Commun, 2017. 8: p. 14532.

96. Shen, M., et al., A novel senolytic drug for pulmonary fibrosis: BTSA1 targets apoptosis of senescent myofibroblasts by activating BAX. Aging Cell, 2024. 23(9): p. e14229.

97. Franzén, L., et al., Mapping spatially resolved transcriptomes in human and mouse pulmonary fibrosis. Nature Genetics, 2024. 56(8): p. 1725–1736.

98. Meng, X.M., D.J. Nikolic-Paterson, and H.Y. Lan, TGF-β: the master regulator of fibrosis. Nat Rev Nephrol, 2016. 12(6): p. 325–38.

99. Feng, Z. and R. Flowerdew. Fuzzy geodemographics: a contribution from fuzzy clustering methods. 1998.

100. Birkin, M. and G. Clarke, *Spatial interaction in geography*. Geography review ; v.4, no.5, 1991, pp.16-24. 1991, Place of publication not identified: [publisher not identified].

101. DeBruine, Z.J., J.A. Pospisilik, and T.J. Triche, Fast and interpretable non-negative matrix factorization for atlas-scale single cell data. bioRxiv, 2024: p. 2021.09.01.458620.

102. Nasution, B.I., R. Kurniawan, and R.E. Caraka, *Nature-Inspired Spatial Clustering [R package naspaclust version 0.2.1]*. 2021, Comprehensive R Archive Network (CRAN).

103. Comber, A., et al., The GWR route map: *a guide to the informed application of Geographically Weighted Regression*. arXiv preprint arXiv:2004.06070, 2020.

104. Bivand, R.S., E. Pebesma, and V. Gómez-Rubio, Applied Spatial Data Analysis with R. 2013, New York, NY, USA: Springer.

105. Bivand, R.S. and D.W.S. Wong, Comparing implementations of global and local indicators of spatial association. TEST, 2018. 27(3): p. 716–748.

106. Cliff, A.D. and J.K. Ord, Spatial autocorrelation. Monographs in spatial and environmental systems analysis ; 5., ed. J.K. Ord. 1973, London: Pion.

107. Cliff, A.D. and J.K. Ord, Spatial processes : models & applications, ed. J.K. Ord and A.D.S.a. Cliff. 1981, London: Pion

