## Supplementary Material for "STExplorer: Navigating the Micro-Geography of Spatial Omics Data"

### Supplementary Methods

#### Spatial transcriptomics analysis of the Prostate dataset

##### Preparation and quality control of sections

10x Visium raw data (H5 files) along with their associated annotations and section images from Erickson et al., 2022[1] were used for the spatial transcriptome analysis of the prostate cancerous sections H1_4, H1_5, H2_1, H2_5 and the benign section V1_2. H5 files were imported as a single Meta Spatial Feature Experiment (msfe) object with the read10xVisiumSFE function from the Bioconductor package SpatialFeatureExperiment[2]. For each section, the following quality control and filtering criteria were applied to remove low-quality spots: **i)** removal of locations with gene expression levels below the 15nth quantile or above the 99nth quantile of all section spots **ii)** removal of locations with library sizes below the 15nth quantile or above the 99nth quantile of all section spots **iii)** removal of locations with mitochondrial content above the 99nth quantile of all section spots. Subsequently, library size factors were calculated for each section with the *computeLibSizeFactors* of STExplorer using default settings, followed by logarithmic transformation with the *normaliseCounts* function of STExplorer. Following log-transformation, genes with low or zero expression were removed with *setQCthresh_LowLogMean*, defining log2=1 as the threshold.

##### NMF analysis

The optimum number of NMF factors was determined using the fgwc_nmfFactorNumber function, using logcounts as the selected assay with default settings, except from k_range which was set to (2,10,1). NMF analysis was performed with the fgwc_nmf function, using the top 10% highly variable genes and selecting the optimum number of factors as ncompoments for each section. Section maps for multiple NMF factors were generated with the plotFGWC_nmfFactorsMap function. Spatial expression of candidate leading genes was visualized as tissue map using the plotGeneExpression function of STExplorer, using logcounts and determining minimum and maximum expression range for each gene manually.

##### Fuzzy Geographical Weighted Clustering

FGWC was performed with the integrated function fgwcSTE of STExplorer, using the classic FGWC algorithm and specifying the distance matrix as Euclidean. The optimum number of FGWC clusters for each section was determined using the fgwc_findOptimumK function of STExplorer with the following arguments: k range between 2 and 10, index type specified as FPC (Fuzzy Partition Coefficient) and knee specified as the elbow method. Winning cluster maps for each section were generated with the plotFGWC_singleMap of STExplorer along with the associated histological annotation using the function plotQC_spotsAnnotation of STExplorer. Multiple cluster maps were generated with the plotFGWC_multiMap function of STExplorer using default settings. Pie-doughnut plots between FGWC and associated histology or between FGWC and NMF factors were generated using the integrated plotFGWC_pie function of STExplorer. Heatmaps with FGWC leading genes were generated with the plotFGWC_nmfFactorsHeatmap of STExplorer, including both the histological annotations and FGWC clusters as annotation columns and independently setting each of them as criteria for ordering the genes on the final heatmap plot.

##### Geographically Weighted Principal Component Analysis

The top 10% of the highly variable genes were isolated for each section with the getTopHighVarGenes function of STExplorer, using an FDR threshold of 0.1. A distance-based neighbor graph was applied to each section, using a knearneigh as type and setting the number of nearest neighbours to six (6). Subsequently, GWPCA analysis along with cross-validation was performed separately for each section using the top 10% highly variable genes with the following arguments: retained components (k) = 20, gaussian kernels, Minkowski distance power (p)=2, fixed bandwidth calculated for each section as 6*average spot diameter. Maps illustrating the locations of single GWPCA leading genes were generated with the plotGWPCA_leadingG function of STExplorer. The percentage of total variation for each section was calculated with the gwpca_PropVar_output of STExplorer for the first ten components and section maps were generated with the plotGWPCA_ptv function of STExplorer.

#### GSEA functional clustering of spatial transcriptome data

For GSEA analysis, the GO BP molecular signatures were first retrieved from MSigDB and then used to associate genes with GO BP terms using the getTerm2Gene function of STExplorer and setting the category as C5 and subcategory as GO:BP. Subsequently, functional clustering was performed with the gwpca_FunctionalClustering function of STExplorer using the following arguments: genes_no=2, NES = 1.5, minGSSize = 5, pAdjustMethod = "fdr", nPermSimple = 10000. Maps with GSEA results were generated with the plotGWPCA_FuncCLust function of STExplorer, setting the minimum cluster count to 8.

#### Isolation of FGWC/NMF and GWPCA meta-signature panels

For composing the FGWC/NMF leading gene meta-signatures for each tissue section, all genes were ranked according to their NMF factor score. The top 10 leading genes were selected based on the following criteria: NMF factor score > average score of the remaining NMF factors (e.g., for selecting NMF1 leaders the criteria is NMF1 factor score > average score of NMF2-5 factors). The composition of the GWPCA meta-signatures relied on the ranking of genes based on component loadings in an annotation-aware manner. More specifically, the average GWPC loading score of all spots that correspond to a certain histological annotation was calculated for each gene and the process was independently repeated for all annotations. Subsequently, all genes were independently ranked based on their loadings and the top 40 or bottom 40 genes were isolated for each histological annotation of each component. The process was repeated for all histological annotations of the first 4 GWPCA components for each tissue section. Subsequent meta-signature analysis focused on the top/bottom 10, 20, 30 or 40 genes of GWPCA1 or the top/bottom 10 genes of GWPCA2-4.

#### Spatial transcriptome analysis with Seurat

Prostate sections were analysed with Seurat according to the standard 10xVisium workflow, using the following pre-processing modifications: **i)** minimum nFeature_Spatial > 15nth quantile and maximum nFeature_Spatial < 99nth quantile **ii)** percent.mt < 15. Normalization was performed with the SCTransform function[3]. Highly variable genes were identified with the FindSpatiallyVariableFeatures Seurat function with moransi as the selection method. Dimensionality reduction was performed both with PCA and UMAP using 30 dimensions of reduction as an input. In each tissue section and for each identified component (n=50 in total), the top and bottom 10 leading genes based on ranked loading scores, were isolated as meta-signatures, composing in total fifty top and fifty bottom 10-gene signature panels.

#### Analysis of scRNA-seq prostate data

For the single-cell analysis of the prostate tumours, data from the Prostate Cell Atlas[4] project were used and analysed with scanpy (https://github.com/scverse/scanpy) in Python. More specifically, the associated scRNA-seq data from this project were imported as H5ad files with the integrated scanpy functions for reading H5 files. QC and preprocessing steps included filtering cells with a mitochondrial reads percentage above 15%. We also filtered out all cells that expressed less than 100 genes. Subsequently, all samples were first normalised using the integrated normalisation function of scanpy and setting the target sum to 10.000 and then were log-transformed. In parallel, an SCVI model was trained using total counts as a continuous covariate and sample name as a categorical covariate. UMAP plots were constructed with the integrated plotting functions for scanpy, using the scvi normalised counts as the selected layer. Dot plots were constructed again with scanpy functions.

##### Analysis of bulk RNA-seq data from TCGA

RNA-seq and clinical data for gastric were downloaded from TCGA with the GDC client. Normal gene expression for gastric biopsies was retrieved from gTEX. For heatmap analysis expression matrixes were cross-normalized, log-transformed and converted to z-scores. Heatmaps were created with the ComplexHeatmap package using the associated clinical information of the tumour samples from TCGA for annotation. ROC analysis was performed as previously described[5], violin plots were created with ggplot2 in R. Forest plots were made with the ggforestplot (https://github.com/NightingaleHealth/ggforestplot) in R. Functional enrichment analysis was performed with the Bioconductor package DOSE[6] in R, dot plots were created with ggplot2. GSEA plots were created with GseaVis (https://github.com/junjunlab/GseaVis)

#### Spatial transcriptomics analysis of the Liver dataset

Human liver spatial transcriptomics data generated on the Visium platform by Guilliams et al., 2022[7] was used for analysis. Two slices were selected: a biopsy with mild pericellular and periportal fibrosis and no steatosis (biopsy ID: JBO018), and one with 70% steatosis and clear pericellular fibrosis (biopsy ID: JBO019), referred to as "mild fibrotic" and "steatotic" respectively. Spots with >15% mitochondrial gene expression and those with the lowest number of features (lowest 5% of spots) and library size (lowest 5% of spots) were filtered out. Additionally, spots where no annotation was provided within the Guilliams et al., 2022 publication[7] were removed. Clustering was performed based on the top 50% highest variable genes with an FDR < 0.05. The number of factors was set at 3 because that resulted in the highest error reduction above noise, as determined by the *fgwc_nmfFactorNumber* function. The optimal number of clusters (k = 5) was selected by the *fgwc_findOptimumK* function. The following gene sets were used for GWPCA functional clustering: Reactome metabolism of lipids [8] and steatosis-, fibrosis-, and NAFLD (Non-Alcoholic Fatty Liver Disease) activity-associated genes from Govaere *et al.*, 2020[9]. Results with a p-value < 0.05 were reported. Selecting the top 10 leading genes (by absolute leading score) for the first three principal components per spot (30 genes per spot), we run a Reactome pathway analysis per spot. We filtered out pathways with less than 2 genes present from the list and for an adjusted p-value of < 0.05. Finally, we calculated the normalised frequencies of each pathway by dividing the number of spots in a cluster the pathway has been enriched by the total number of spots in the cluster.

#### Spatial transcriptomics analysis of the Lung dataset

##### Preparation and quality control of sections

Data were downloaded from Franzén L et al., Nat Genetics, 2024[10]. The published spatial sequencing dataset of lung tissues was derived from 4 healthy individuals (healthy controls; HC) and 4 IPF patients. We used 16 lung slices with different grades-extent of fibrosis (healthy, FS1-mild fibrosis, FS2-moderate fibrosis, FS3-severe fibrosis, by histological inspection). The specific sample IDs used are: V19S23-092-A1, V19S23-092-B1, V19S23-092-C1, V19S23-092-D1, V10T31-015-A1, V10T31-051-B1, V10T31-051-C1, V10T31-051-D1, V10T03-280-A1, V10T03-280-B1, V10T03-280-C1, V10T03-280-D1, V10T03-281-

A1, V10T03-281-B1, V10T03-281-C1, and V10T03-281-D1. More information about the slice metadata can be found in the original publication and on GitHub with the code for this publication's analysis. The data Quality Control and filtering were applied in both spots and genes. The settings were a minimum of 350 UMIs per spot, 100 UMIs per gene, and 100 genes per spot. Additional filtering excluded spots with more than 30% mitochondria and/or haemoglobin gene expression. Gene information was retrieved via biomaRt[11] and used to select for 'protein-coding', 'IG' (immunoglobulin) and 'TR' (T cell receptor) gene biotypes, as well as to flag X and Y chromosome genes for removal to avoid sex biases. After removing low-quality spots and genes and before normalisation, all 16 samples were added to a Seurat object to perform normalisation and scaling with *SCTransform* (Seurat package). For the normalisation, sample ID and donor were specified as variables to regress out to remove major effects of technical and interindividual differences.

All thresholds for filtering were set based on an initial examination of the raw data to exclude low-quality spots and genes with low expression.

#### Score calculation

To detect senescent cells, we employed the SenMayo score, a well-established gene set that identifies senescent cells and predicts senescence-associated pathways across tissues (Saul et al., Nat Commun, 2022). For fibrosis detection, we derived the Extracellular Matrix (ECM) score using genes from the Molecular Signatures Database (MSigDB) Gene Ontology Cellular Component GO_EXTRACELLULAR_MATRIX (https://www.gsea-msigdb.org/gsea/msigdb/human/geneset/GO_EXTRACELLULAR_MATRIX). The epithelial-to-mesenchymal transition (EMT) score was generated using genes from the MSigDB Hallmark gene set EPITHELIAL_MESENCHYMAL_TRANSITION (https://www.gsea-msigdb.org/gsea/msigdb/human/geneset/HALLMARK_EPITHELIAL_MESENCHYMAL_TRANSITION). Lastly, the TGF-β score was calculated based on the gene set identified by Ma et al., 2024, focusing on genes induced in fibroblasts following TGF-β stimulation. All scores were computed using the *AddModuleScore* function from Seurat with default parameters applied to the specified gene lists. The 75th percentile (top 25% of values) was used as the cut-off to distinguish areas with high versus low senescence, fibrosis, EMT, and TGFB scores.

#### Geographically Weighted Regression analysis

We performed Geographically Weighted Regression (GWR) analysis using the *gwrSTE* function from STExplorer. As input we used the *"senMayo~ecm"* formula for regression between senescence and fibrosis ('formula' argument), the basic GWR method ('gwr_method' argument), 3 times the spot diameter as neighbourhood-setting bandwidth ('bw' argument), and an exponential kernel to weigh the distances in space ('kernel' argument). We set the GWR $\beta_1$ coefficient cut-off to 0.5 to capture areas with high positive correlation between senescence and fibrosis. The cell type densities used in the analysis were taken from Franzén et. al.[10].
Spatial Autocorrelation statistics were calculated for the SenMayo score using *getisLocalGPerm* function from STExplorer, providing the SenMayo scores per spot as input with default parameters.

#### Spatial transcriptomics analysis of the developmental Eye dataset

An eight post-conception week (PCW) developing eye sample (8PCW_D) generated using 10X Visium from the published dataset[12] was analysed using STExplorer version 0.1.0 (commit 398f149). Before analysis, the sample was loaded into a SpatialFeatureExperiment v1.4.0 object in R 4.3.2. Spots with > 15% mitochondrial gene expression, <500 genes and a library size of <500 or >32000 were removed from further downstream analysis. Ground-truth annotations were taken from the published dataset

[12]. Spots annotated with ‘Vitreous’ or ‘RBC’ were also removed at this stage. As the vitreous consists of an extracellular matrix that presumably contains no RNA transcripts, any counts were presumed to be because of contamination. Spots annotated as red blood cells (RBCs) were removed as the analysis was dominated by haem genes downstream.

The optimum number of NMF factors was determined using the *fgwc_nmfFactorNumber* function from STExplorer. Five (5) NMF factors were used. The value of *k* for the number of clusters was determined using the STExplorer function *fgwc_findOptimumK*, with ‘composite’ passed to the *index_type* argument. Eight (8) was determined to be the optimum value of *k*. All other settings were the default settings as described in the STExplorer vignette. Highly variable genes (HVGs) were identified using the *modelGeneVar* and *getTopHVGs* functions from scran 1.30.2[13]. One hundred and fifty (150) genes were identified as highly variable. Following filtering for only protein-coding genes, 90 genes were identified as highly variable. Fuzzy geographical weighted clustering (FGWC) was run using the *fgwcSTE* function passing as *k=8* the number of clusters and retaining the rest parameters at default.

### Supplementary Figures

#### Supplementary Figure 1

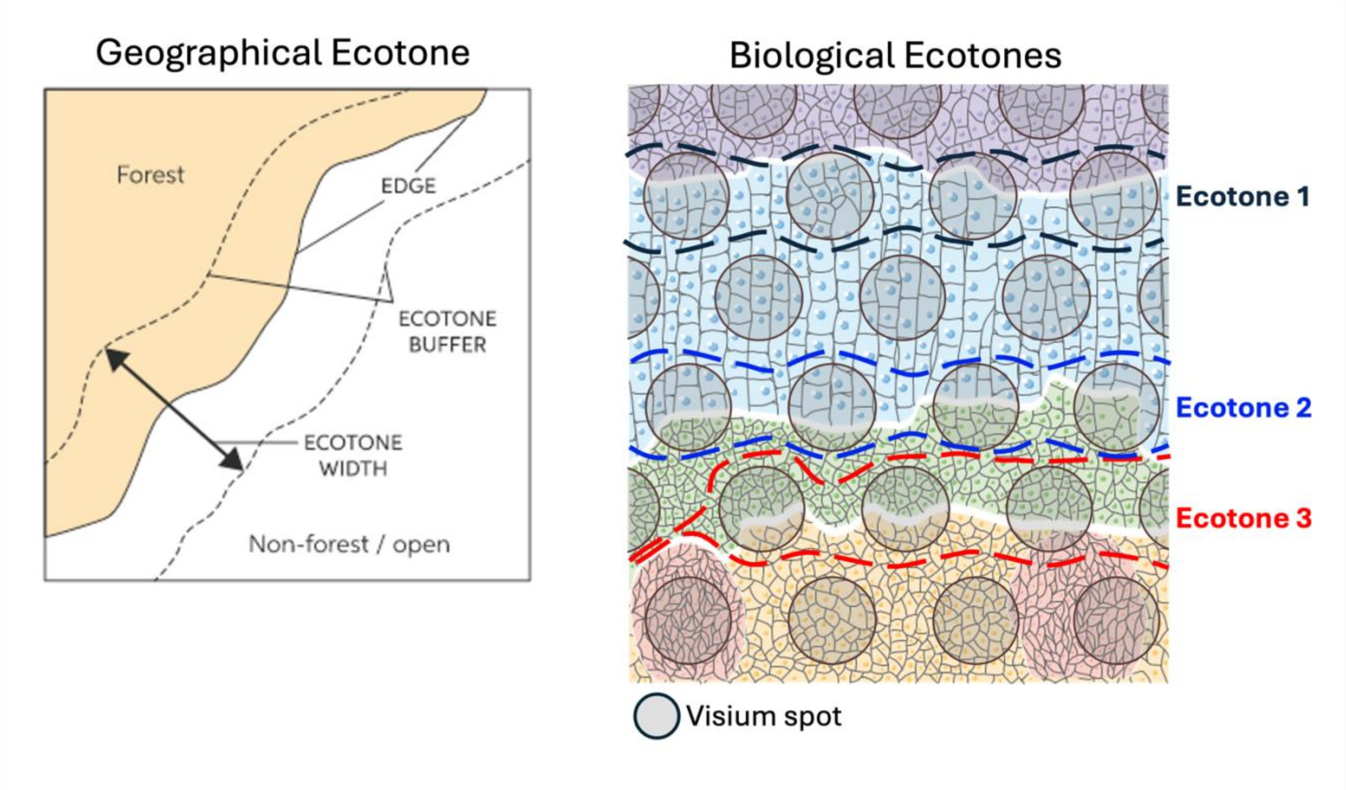

**Supp. Figure 1 Spatial ecotones.** The left panel is an example of a geographical ecotone as found in nature. An ecotone is a zone that overlaps the boundary between the forest and a plain field. This zone is usually defined by a gradient where the forest is gradually thinning until it "becomes" a plain field. Not all forests have ecotones covering their edges and not all ecotones have the same width. The right panel is showcasing the idea of Biological ecotones. In this case the ecotones are introduced artificially due to the platform used for the experiment (in this case Visium). For example, a Visium platform with a fixed spots array generates ecotones by having spots overlapping two or more regions on the tissue. Another example of ecotones can be found in tumour samples where we might encounter cells bordering tumorous areas that undergo molecular changes in different levels, thus not yet fully transitioning from healthy to tumorous.

#### Supplementary Figure 2

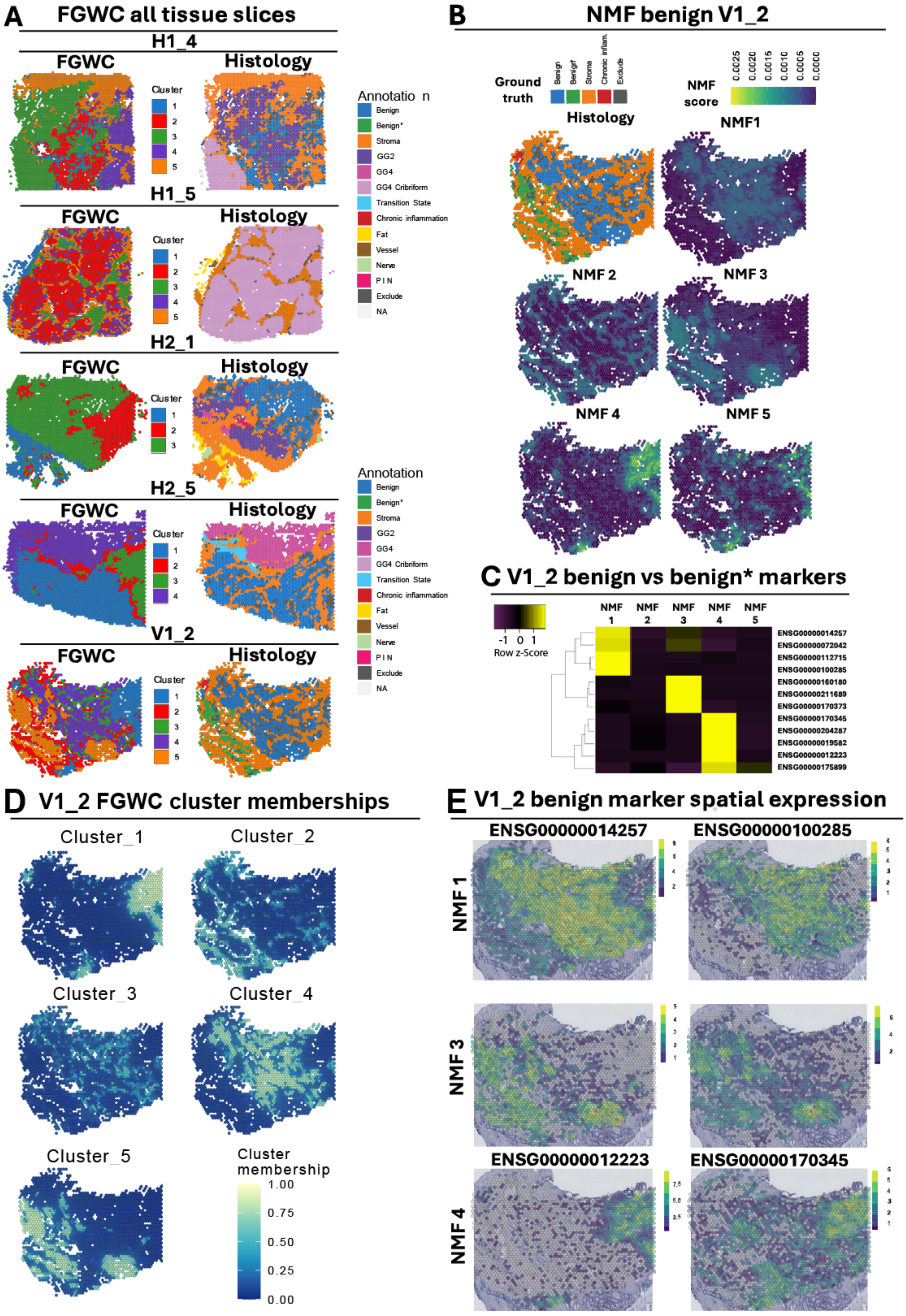

**Supplementary Figure 2 FGWC analysis of prostate section V1_2.**

**A.** Juxtaposition of FGWC cluster distribution against histological annotation for all prostate tissue sections **B.** NMF factor identification for benign section V1_2. Note that NMF3 is mostly enriched in benign*, which corresponds to genomically amplified benign epithelium. **C.** Heatmap illustrating the score of the top leading genes per NMF factor for V1_2. **D.** Multi-cluster membership maps for the V1_2 slice. Each spot on the map has a membership % in each of the 5 clusters. **E.** Representative examples of spatial gene expression for selected NMF factors 1 (mostly benign epithelium), 3 (mostly benign* epithelium) and 4 (mostly stroma).

**References**

ENSG00000014257 ⇔ ACP3/ PPAP[14, 15]
ENSG00000100285 ⇔ NEFH[16, 17]
ENSG00000211689 ⇔ TRGC1[18, 19]
ENSG00000160180 ⇔ TFF3[20, 21]
ENSG00000012223 ⇔ LTF[22, 23]
ENSG00000170345 ⇔ FOS[24]

### Supplementary Figure 3

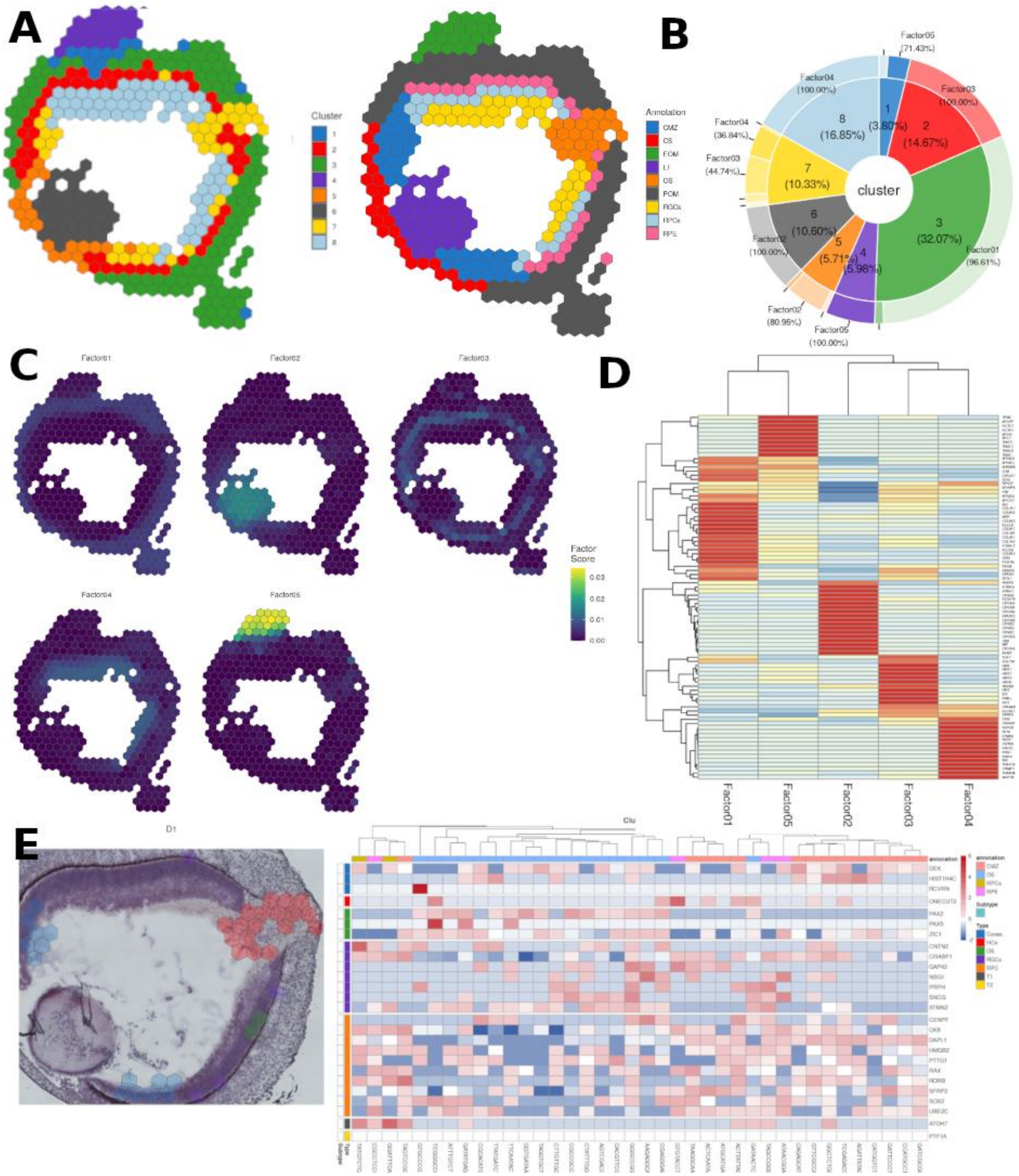

**Supplementary Figure 3 FGWC analysis of the developmental eye**

**A.** FGWC identified 8 clusters, which broadly corresponds to the ground truth annotation. The ciliary margin zone (CMZ) is covered by cluster 2, 5 and 7. The extra-ocular muscle (EOM) is captured faithfully by cluster 5. The peri-ocular mesenchyme is covered by clusters 1 and 3. The corneal stroma corresponds to cluster 5. Lens fibre (LF) is recovered completely by cluster 6. The optic stalk (OS) is covered by 2 clusters: cluster 2 and cluster 7. The retinal ganglion cell layer is not recovered, but forms part of cluster 8. Similarly, the retinal progenitor cell (RPC) layer is not recovered and forms part of cluster 8. The retinal progenitor epithelium is recovered by cluster 2, though cluster 2 also covers the rear of the ciliary margin zone. **B.** Pie-doughnut chart showcasing the combination of NMF factors leading to each of the 5 clusters. Factor03 and 04 contribute the most to cluster 7. **C.** NMF Factor maps revealing the spatial organisation of each factor. **D.** The NMF factor score of each gene display their contribution to each factor. The top genes (excluding haemoglobin, mitochondrial and ribosomal genes in Factor03 are: PMEL, VIM, ID3, DCT, SLC2A1, SPARC, CRABP1 and CKB. The top genes in Factor04 (again excluding ribosomal, haemoglobin and mitochondrial genes) are: CRABP1, TUBA1A, TUBB2B, CKB, STMN2, MAP1B, VIM, CNTN2 and CRMP1. Noteworthy in Factor03 are the genes PMEL and DCT which are known markers for the iris pigment epithelium, ID3 and CKB which are known markers for RPCs and VIM which is an established marker for muller glia cells. In Factor04

CRMP1 belongs to the CRMP family of genes that have a role in establishing neuronal polarity, and CNTN2, CRABP1 and STMN2 which are known markers for RGCs. **E.** Cluster 7 location coloured by ground truth alongside a heatmap of marker gene expression per spot organised by ground truth. Cluster 7 could potentially be of biological significance as it covers the OS and CMZ, which are potential sources of RPCs during retinal development [12]. A similar cluster was identified in the original publication; however, this was achieved by plotting early RPC markers and was not identified from graph-based clustering.

Supplementary Figure 4

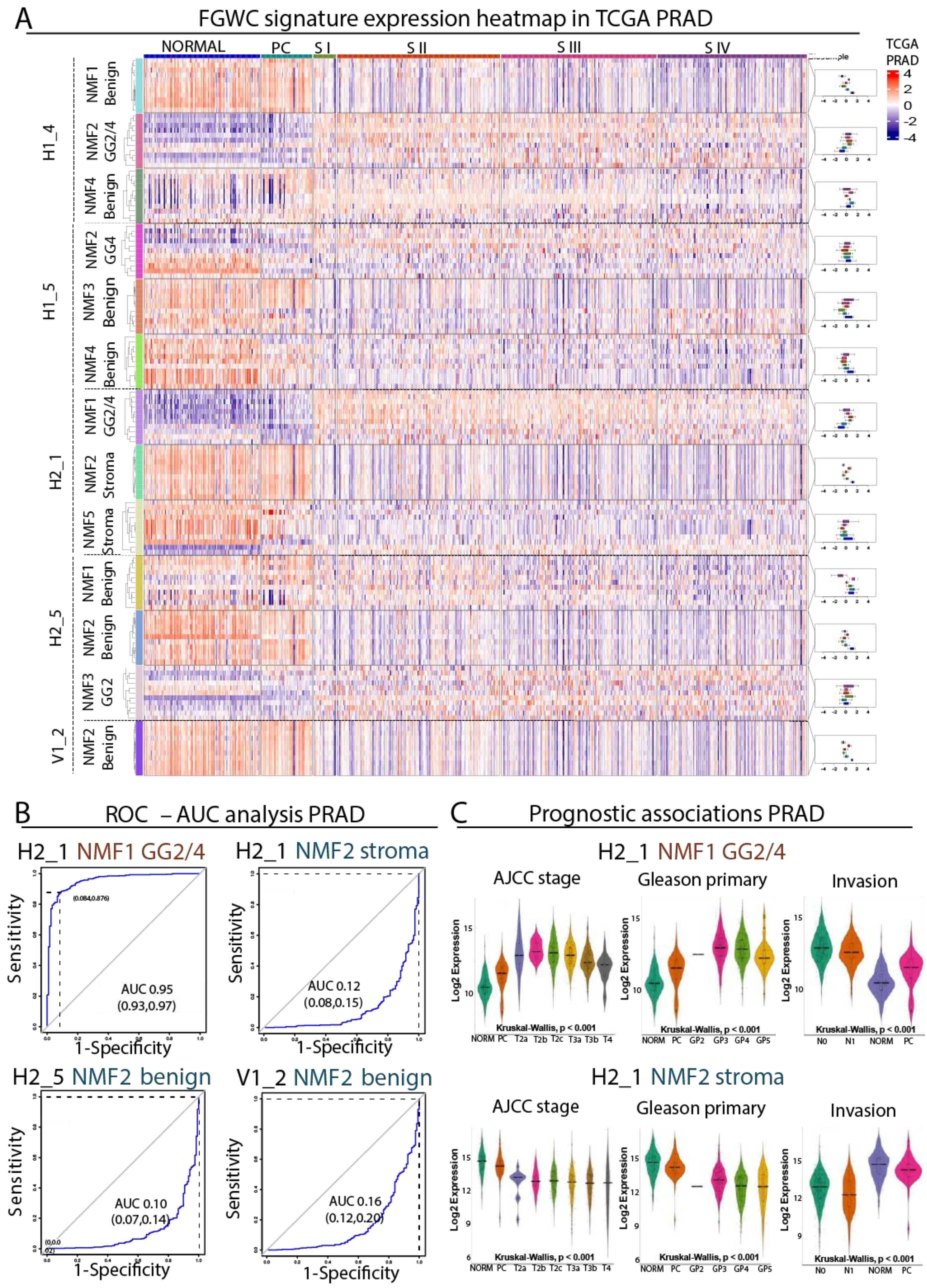

**Supplementary Figure 4 FGWC/NMF meta-signature performance in TCGA-PRAD bulk RNA-seq. A.** Heatmap summarizing the expression (z-scores of log2 expression) of all NMF metagene signatures in staged gTEX and TCGA-PRAD bulk RNA-seq data. Biopsies are annotated with the colored bar at the top (PC: paracancerous, S I-IV: Stage I-IV). **B.** Representative ROC-AUC analysis for prostate tumor diagnosis of two cancerous (top panels, referring to prostate sections H1_4 and H2_1) and two benign (bottom panels, referring to prostate sections H2_5 and V1_2) NMF signatures. Average AUC with 95% CI are shown in parenthesis along with sensitivity and specificity performance. **C.** Violin plots illustrating the expression of two NMF metagene signatures in TCGA-PRAD biopsies stratified according to AJCC tumor stage (left), primary Gleason score (center) or AJCC lymph node invasion (right). Top panels represent a cancerous signature from H1_4 and the bottom panels correspond to the benign signature from V1_2 prostate section. Kruskal-Wallis corresponds to statistical comparisons against the expression in the normal biopsies.

#### Supplementary Figure 5

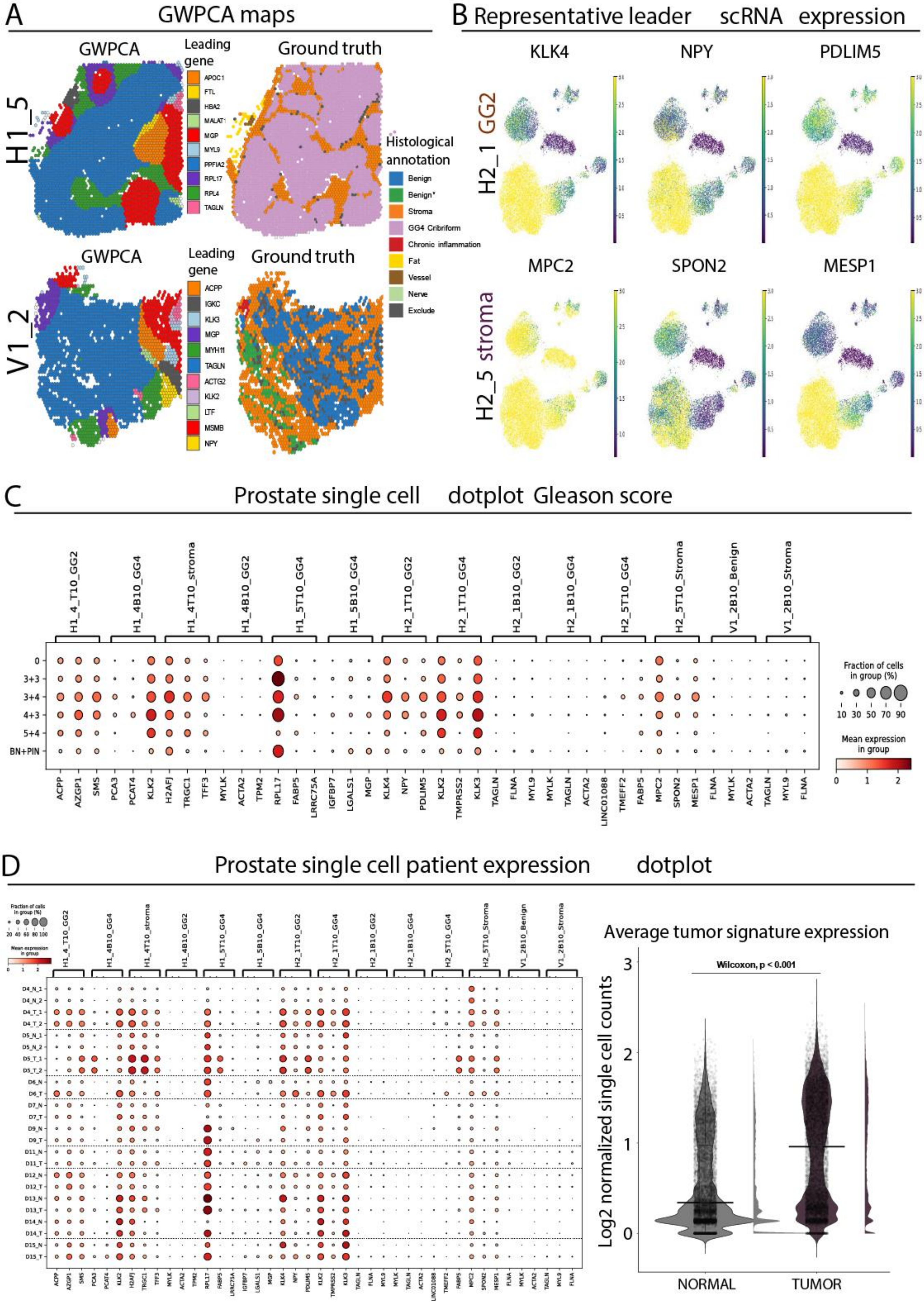

**Supplementary Figure 5 Analysis of GWPCA metagene signature expression in prostate scRNA-seq data A.** GWPCA maps (left) with histological annotations (right) for the cancerous prostate section H1_5 (top) and the benign section V1_2 (bottom). **B.** scRNA-seq uMAPs summarizing the expression of three representative leading genes from sections H2_1 (top, derived from GG2 spots) and H2_5 (bottom, derived from tumor surrounding stroma). **C.** Dot plot overview of three representative leaders for indicative GWPCA meta-signatures from all sections across the scRNA-seq panel, after stratification according to primary and secondary gleason score. **D.** Same as (C) but samples are stratified into tumor (T) and normal (N) for each patient ID. Horizontal dashed lines separate the different patients. Violin plot summarizes the expression of all cancerous signatures in normal vs tumor cells. Pvalue corresponds to statistical significance according to Wilcoxon test.

Supplementary Figure 6

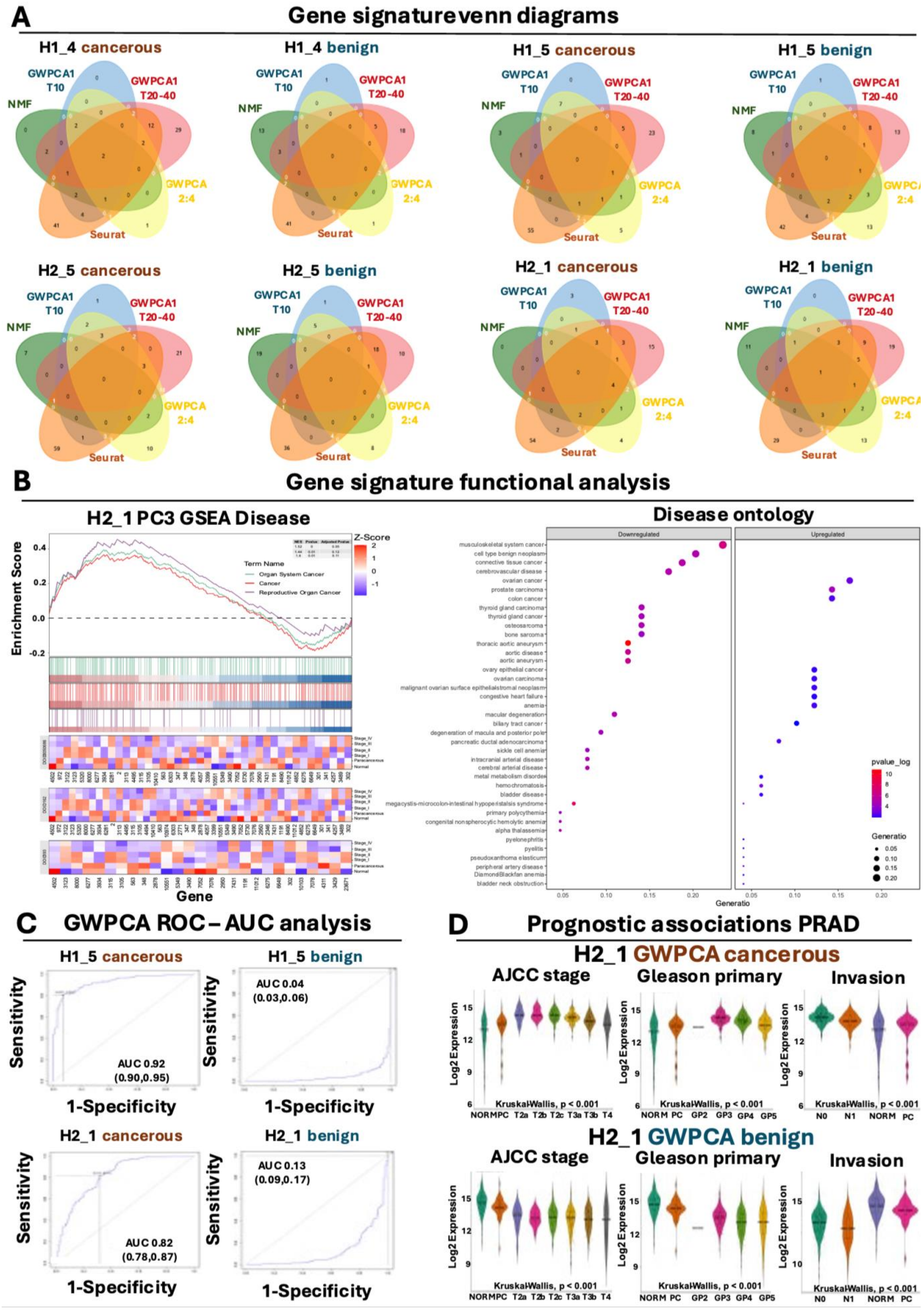

**Supplementary Figure 6 FGWC and GWPCA meta-signature analysis in TCGA PRAD bulk RNA-seq data. A.** Venn diagrams comparing meta-signature composition for all cancerous sections. NMF corresponds to all unique leaders from NMF factors, GWPCA1 T10 corresponds to the top 10 unique leaders of GWPCA1, GWPCA T20-40 corresponds to the remaining 20 to 40 top unique leaders from GWPCA1, GWPCA2:4 correspond to the top 10 unique leaders of GWPCA components 2,3 and 4, while Seurat corresponds to the top 10 unique leaders of uMAP clusters 1 to 10. **B.** Functional analysis of GWPCA metagene signature for the H2_1 section. The left panel shows GSEA disease enrichment for Organ System Cancer (turquoise), cancer (red) and reproductive organ cancer (purple) categories for genes that are stratified according to GWPCA3 loadings. Bottom heatmaps demonstrate representative gene expression from each category in staged gTEX/TCGA-PRAD biopsies. The panel on the right summarizes disease enrichment of GWPCA metagene signatures, separated according to their expression in TCGA-PRAD data. Downregulated refers to leaders that reduce their expression in prostate cancerous tissues, while upregulated refers to leaders that are overexpressed in prostate tumors. **C.** Representative ROC-AUC analysis for prostate tumor diagnosis of one cancerous (left) and one benign (right) GWPCA meta-signature from sections H1_5 (top) and H2_1 (bottom). Average AUC with 95% CI are shown in parenthesis along with sensitivity and specificity performance. **D.** Violin plots illustrating the expression of the GWPCA1 cancerous (top) and benign (bottom) meta-signatures from section H2-1 in TCGA-PRAD biopsies stratified according to AJCC tumor stage (left), primary Gleason score (center) or AJCC lymph node invasion (right). Kruskal-Wallis corresponds to statistical comparisons against the expression in the normal biopsies.

### Supplementary Figure 7

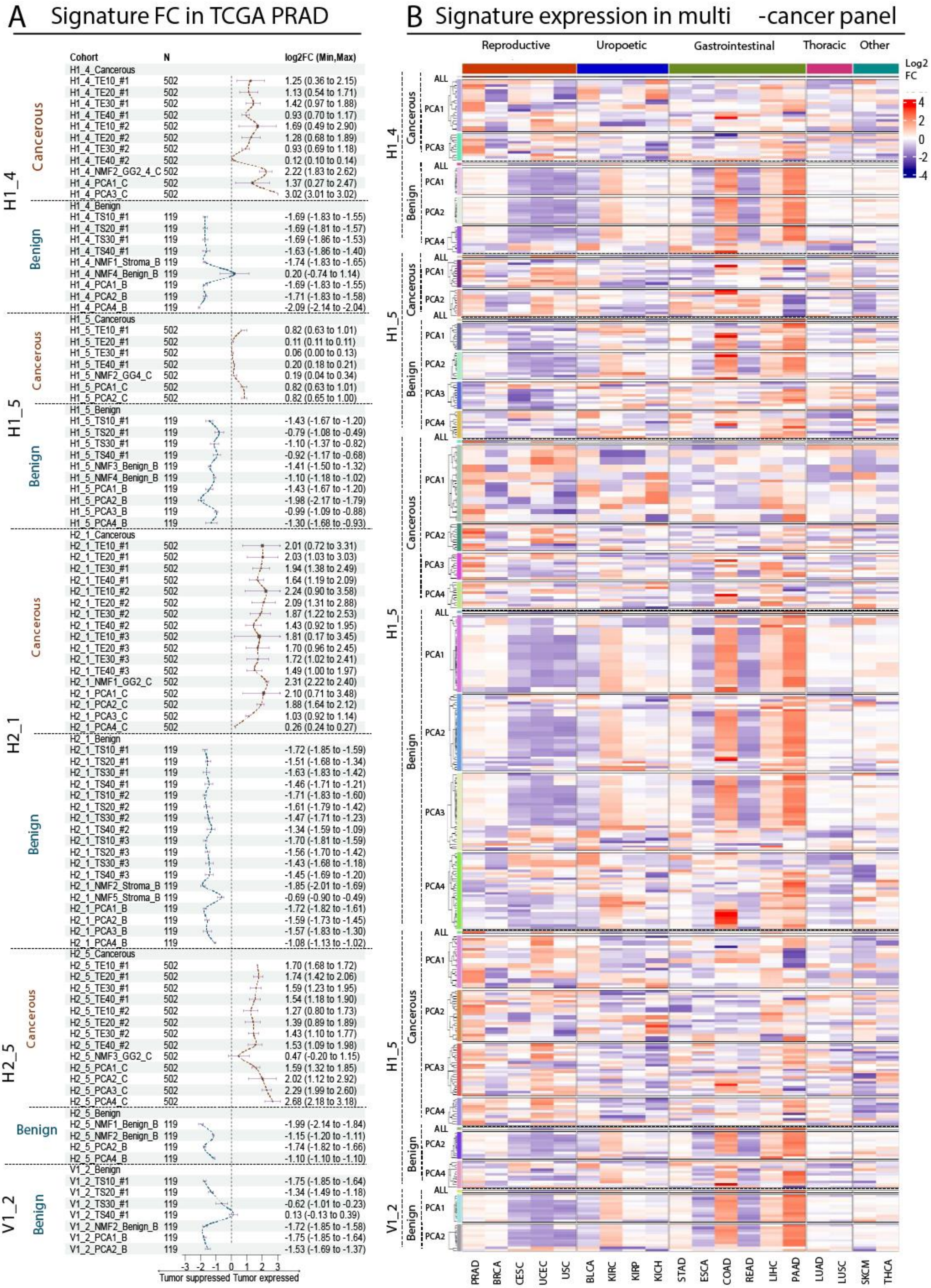

**Supplementary Figure 7 FGWC and GWPCA meta-signature analysis across multiple TCGA cancer panels. A.** Forest plot summarizing the fold change of all FGWC and GWPCA meta-signatures in TCGA-PRAD data. Expression change is shown as log2FC along with min and max changes in parenthesis. The vertical dashed line corresponds to absence of differential expression, red curves correspond to upregulation in prostate tumor biopsies while blue curves correspond to upregulation in normal (GTEx) prostate biopsies. TS: Tumor-suppressed, TE: Tumor-expressed, TE10-40: GWPCA1 top 10-40 genes, PCA1-4: GWPCA1-4 top10 genes. **B.** Heatmap summarizing the fold change of representative GWPCA signatures across a multi-cancer TCGA panel, shown as z-scores. Red corresponds to average log2 fold chance increase, blue corresponds to average log2 fold change decrease of each signature in the corresponding cancer type. Tumor types are organized according to system origins as indicated with the top-colored bar. Cancer names are shown at the bottom and correspond to TCGA abbreviations. PCA1-4: GWPCA1-4 top10 genes.

#### Supplementary Figure 8

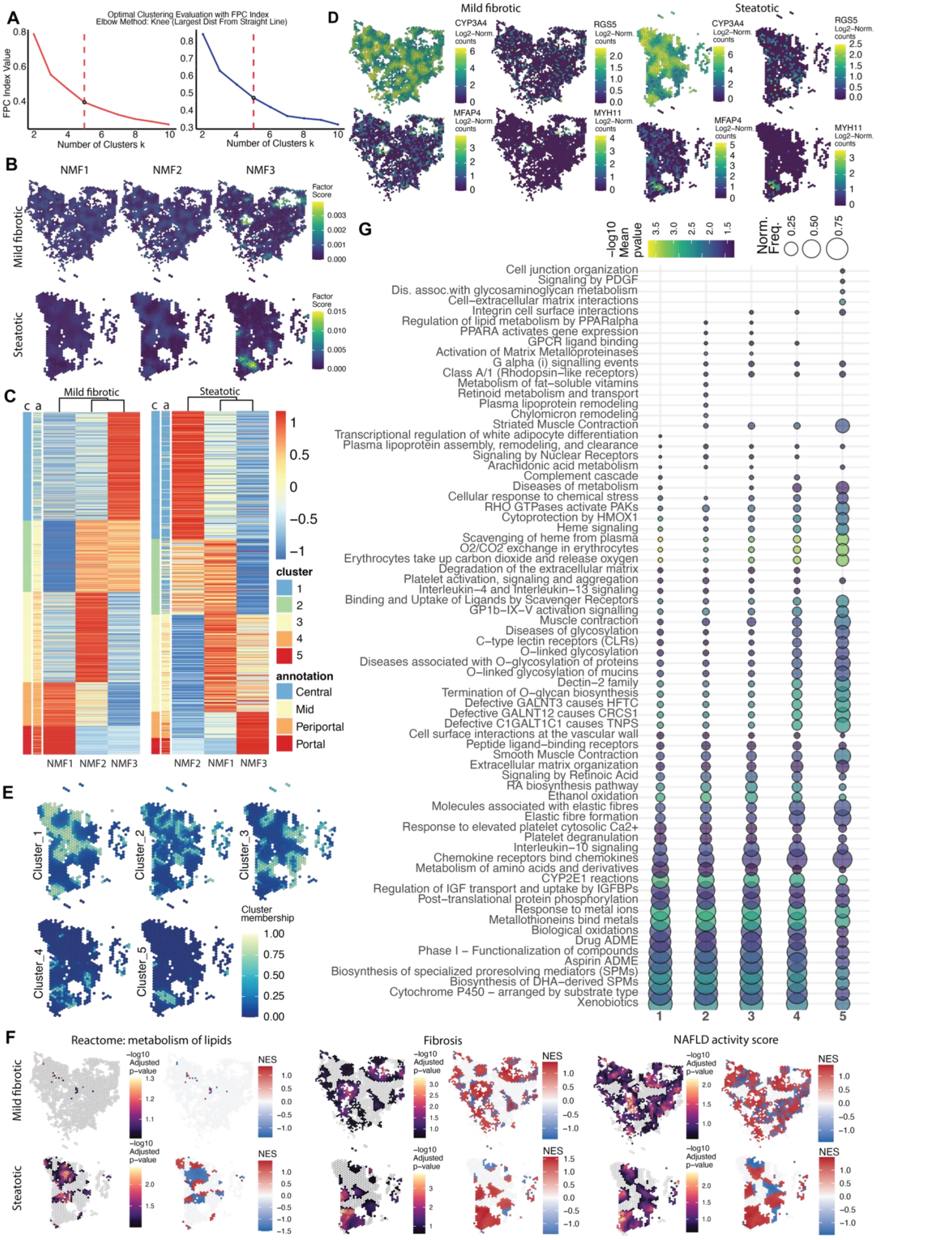

**Supplementary Figure 8 FGWC and GWPCA analysis across liver samples. A.** Fuzzy Partition Coefficient elbow plots for the identification of the optimum number of clusters (k) for the Mild fibrotic (left) and Steatotic (right) samples. **B.** NMF factor score maps visualizing the spatial patterns of the NMF factors the combination of whom gives rise to the 5 clusters. **C.** NMF factor heatmaps ordered by winning cluster showcasing the NMF factor patterns. **D.** Spatial expression maps of key liver genes related to the tissue section biology. **E.** Membership percentage maps of the Steatotic sample showcasing putative ecotones between clusters 2 and 3. **F.** Adjusted p-values and Normalised Enrichment Scores for the pathways used in the Functional Clustering. **G.** Full table of the Reactome pathway enrichment of the Mild-fibrotic sample using the top 10 leading genes from the first three principal components (30 genes in total) in each spot.

### Supplementary Table 1

| Sample | Platform | # of HVGs | Bandwidth | Iteration locations | Kernel | k | Minkowski distance | Score calculation | Robust GWPCA | Cross-Validation | Elapsed time (mins) | PC1 % | PC2 % | PC3 % | Total PC1-3 % |
|---|---|---|---|---|---|---|---|---|---|---|---|---|---|---|---|
| H1_4 | Windows with 7/16 dedicated cores / PSOCK | 1770 (top 50%) | 680.0001 | 3930 | gaussian | 20 | Euclidean | NO | NO | YES | 1188.591 | 2.783 | 1.089 | 0.724 | 4.596 |
| H1_4 | Windows with 7/16 dedicated cores / PSOCK | 354 (top 10%) | 680.0001 | 3930 | gaussian | 20 | Euclidean | NO | NO | YES | 29.474 | 10.394 | 4.535 | 3.28 | 18.209 |
| H1_5 | Windows with 7/16 dedicated cores / PSOCK | 228 (top 10%) | 566.6403 | 3707 | gaussian | 20 | Euclidean | NO | NO | YES | 9.365 | 10.07 | 6.53 | 2 | 18.6 |
| V1_2 | Windows with 7/16 dedicated cores / PSOCK | 337 (top 10%) | 679.8512 | 2621 | gaussian | 20 | Euclidean | NO | NO | YES | 10.394 | 21.62 | 8.73 | 4.37 | 34.72 |
| H1_4 | Windows with 7/16 dedicated cores / PSOCK | 708 (top 20%) | 680.0001 | 3930 | gaussian | 20 | Euclidean | NO | NO | YES | 139.362 | 6.038 | 2.505 | 1.731 | 10.274 |
| H1_5 | Windows with 7/16 dedicated cores / PSOCK | 455 (top 20%) | 566.6403 | 3707 | gaussian | 20 | Euclidean | NO | NO | YES | 44.685 | 6.142 | 3.687 | 1.088 | 10.917 |
| V1_2 | Windows with 7/16 dedicated cores / PSOCK | 674 (top 20%) | 679.8512 | 2621 | gaussian | 20 | Euclidean | NO | NO | YES | 50.61 | 13.74 | 5.38 | 3.41 | 22.53 |
| H1_4 | Linux with 16/32 dedicated cores / FORK | 1770 (top 50%) | 680.0001 | 3930 | gaussian | 20 | Euclidean | NO | NO | YES | 995.906 | 2.783 | 1.089 | 0.724 | 4.596 |
| H1_4 | Linux with 16/32 dedicated cores / FORK | 354 (top 10%) | 680.0001 | 3930 | gaussian | 20 | Euclidean | NO | NO | YES | 17.387 | 10.394 | 4.535 | 3.28 | 18.209 |
| H1_5 | Linux with 16/32 dedicated cores / FORK | 228 (top 10%) | 566.6403 | 3707 | gaussian | 20 | Euclidean | NO | NO | YES | 4.705 | 10.07 | 6.53 | 2 | 18.6 |
| V1_2 | Linux with 16/32 dedicated cores / FORK | 337 (top 10%) | 679.8512 | 2621 | gaussian | 20 | Euclidean | NO | NO | YES | 4.74 | 21.62 | 8.73 | 4.37 | 34.72 |
| H1_4 | Linux with 16/32 dedicated cores / FORK | 708 (top 20%) | 680.0001 | 3930 | gaussian | 20 | Euclidean | NO | NO | YES | 102.733 | 6.038 | 2.505 | 1.731 | 10.274 |
| H1_5 | Linux with 16/32 dedicated cores / FORK | 455 (top 20%) | 566.6403 | 3707 | gaussian | 20 | Euclidean | NO | NO | YES | 27.02 | 6.142 | 3.687 | 1.088 | 10.917 |
| V1_2 | Linux with 16/32 dedicated cores / FORK | 674 (top 20%) | 679.8512 | 2621 | gaussian | 20 | Euclidean | NO | NO | YES | 33.063 | 13.74 | 5.38 | 3.41 | 22.53 |

**Supplementary Table 1 GWPCA computational performance metrics.**

**Sample:** Prostate cancer sample ID, **Platform:** machine specs used, **#ofHVGs:** number of High Variable Genes (HVGs) fed to GWPCA, **Bandwidth:** the selected bandwidth for GWPCA, **Iteration locations:** locations included in the sample that GWPCA iterated over, **Kernel:** kernel type used in GWPCA, **k:** number of retained principal components, **Minkowski distance:** distance metric used – Euclidean equates to physical distance between locations, **Score calculation:** was score stored, **Robust GWPCA:** was robust GWPCA run, **Cross-Validation:** was cross-validation of local PCA results performed, **Elapsed time (mins):** time elapsed to complete GWPCA in minutes.

### References

1. Erickson, A., et al., *Spatially resolved clonal copy number alterations in benign and malignant tissue*. Nature, 2022. **608**(7922): p. 360-367.
2. Moses, L., et al., *Voyager: exploratory single-cell genomics data analysis with geospatial statistics*. bioRxiv, 2023: p. 2023.07.20.549945.
3. Hafemeister, C. and R. Satija, *Normalization and variance stabilization of single-cell RNA-seq data using regularized negative binomial regression*. Genome Biology, 2019. **20**(1): p. 296.
4. Tuong, Z.K., et al., *Resolving the immune landscape of human prostate at a single-cell level in health and cancer*. Cell Rep, 2021. **37**(12): p. 110132.
5. Samara, M., et al., *Characterization of a miRNA Signature with Enhanced Diagnostic and Prognostic Power for Patients with Bladder Carcinoma*. International Journal of Molecular Sciences, 2023. **24**(22): p. 16243.
6. Yu, G., et al., *DOSE: an R/Bioconductor package for disease ontology semantic and enrichment analysis*. Bioinformatics, 2015. **31**(4): p. 608-9.
7. Guilliams, M., et al., *Spatial proteogenomics reveals distinct and evolutionarily conserved hepatic macrophage niches*. Cell, 2022. **185**(2): p. 379-396.e38.
8. Reactome, r. *Metabolism of lipids*. 20-12-2024]; Available from: https://reactome.org/PathwayBrowser/#/R-HSA-556833.
9. Govaere, O., et al., *Transcriptomic profiling across the nonalcoholic fatty liver disease spectrum reveals gene signatures for steatohepatitis and fibrosis*. Sci Transl Med, 2020. **12**(572).
10. Franzén, L., et al., *Mapping spatially resolved transcriptomes in human and mouse pulmonary fibrosis*. Nature Genetics, 2024. **56**(8): p. 1725-1736.
11. Smedley, D., et al., *BioMart--biological queries made easy*. BMC Genomics, 2009. **10**: p. 22.
12. Dorgau, B., et al., *Single-cell analyses reveal transient retinal progenitor cells in the ciliary margin of developing human retina*. Nature Communications, 2024. **15**(1): p. 3567.
13. Lun, A.T., D.J. McCarthy, and J.C. Marioni, *A step-by-step workflow for low-level analysis of single-cell RNA-seq data with Bioconductor*. F1000Res, 2016. **5**: p. 2122.
14. Xu, H., et al., *Prostatic Acid Phosphatase (PAP) Predicts Prostate Cancer Progress in a Population-Based Study: The Renewal of PAP?* Dis Markers, 2019. **2019**: p. 7090545.
15. Han, H., et al., *Prostate epithelial genes define therapy-relevant prostate cancer molecular subtype*. Prostate Cancer Prostatic Dis, 2021. **24**(4): p. 1080-1092.
16. Su, Z., G. Wang, and L. Li, *CHRDL1, NEFH, TAGLN and SYNM as novel diagnostic biomarkers of benign prostatic hyperplasia and prostate cancer*. Cancer Biomark, 2023. **38**(2): p. 143-159.
17. Peng, Q., et al., *PiRNA-4447944 promotes castration-resistant growth and metastasis of prostate cancer by inhibiting NEFH expression through forming the piRNA-4447944-PIWIL2-NEFH complex*. Int J Biol Sci, 2024. **20**(9): p. 3638-3655.
18. Cerapio, J.P., et al., *Phased differentiation of γδ T and T CD8 tumor-infiltrating lymphocytes revealed by single-cell transcriptomics of human cancers*. Oncoimmunology, 2021. **10**(1): p. 1939518.
19. Chelebian, E., et al., *Morphological Features Extracted by AI Associated with Spatial Transcriptomics in Prostate Cancer*. Cancers (Basel), 2021. **13**(19).
20. Liu, J., et al., *Overexpression of TFF3 is involved in prostate carcinogenesis via blocking mitochondria-mediated apoptosis*. Exp Mol Med, 2018. **50**(8): p. 1-11.
21. Timofte, A.D., et al., *HOXB13 and TFF3 can contribute to the prognostic stratification of prostate adenocarcinoma*. Rom J Morphol Embryol, 2021. **62**(1): p. 41-52.
22. Zhao, Q., Y. Cheng, and Y. Xiong, *LTF Regulates the Immune Microenvironment of Prostate Cancer Through JAK/STAT3 Pathway*. Front Oncol, 2021. **11**: p. 692117.
23. Song, H., et al., *Single-cell analysis of human primary prostate cancer reveals the heterogeneity of tumor-associated epithelial cell states*. Nat Commun, 2022. **13**(1): p. 141.
24. Riedel, M., et al., *In vivo CRISPR inactivation of Fos promotes prostate cancer progression by altering the associated AP-1 subunit Jun*. Oncogene, 2021. **40**(13): p. 2437-2447.
